# Coupling chromatin structure and dynamics by live super-resolution imaging

**DOI:** 10.1101/777482

**Authors:** R. Barth, K. Bystricky, H. A. Shaban

## Abstract

Chromatin conformation regulates gene expression and thus constant remodeling of chromatin structure is essential to guarantee proper cell function. To gain insight into the spatio-temporal organization of the genome, we employ high-density photo-activated localization microscopy and deep learning to obtain temporally resolved super-resolution images of chromatin in living cells. In combination with high-resolution dense motion reconstruction, we confirm the existence of elongated ~ 45 to 90 nm wide chromatin ‘blobs’. A computational chromatin model suggests that these blobs are dynamically associating chromatin fragments in close physical and genomic proximity and adopt TAD-like interactions in the time-average limit. Experimentally, we found that chromatin exhibits a spatio-temporal correlation over ~ 4 μm in space and tens of seconds in time, while chromatin dynamics are correlated over ~ 6 μm and last 40 s. Notably, chromatin structure and dynamics are closely related, which may constitute a mechanism to grant access to regions with high local chromatin concentration.

## Introduction

The three-dimensional organization of the eukaryotic genome plays a central role in gene regulation (*1*). Its spatial organization has been prominently characterized by molecular and cellular approaches including high-throughput chromosome conformation capture (Hi-C) (*2*) and fluorescent in situ hybridization (FISH) (*3*). Topologically associated domains (TADs), genomic regions that display a high degree of interaction, were revealed and found to be a key architectural feature (*4*). Direct 3D localization microscopy of the chromatin fiber at the nanoscale (*5*) confirmed the presence of TADs in single cells but also, among others, revealed great structural variation of chromatin architecture (*3*). To comprehensively resolve the spatial heterogeneity of chromatin, super-resolution microscopy must be employed. Previous work showed that nucleosomes are distributed as segregated, nanometer-sized accumulations throughout the nucleus (*6*–*8*) and that the epigenetic state of a locus has a large impact on its folding (*9*, *10*). However, to resolve the fine structure of chromatin, high labeling densities, long acquisition times and, often, cell fixation are required. This precludes capturing dynamic processes of chromatin in single live cells, yet chromatin moves at different spatial and temporal scales.

The first efforts to relate chromatin organization and its dynamics were made using a combination of Photo-activated Localization Microscopy (PALM) and tracking of single nucleosomes (*11*). It could be shown that nucleosomes mostly move coherently with their underlying domains, in accordance with conventional microscopy data (*12*); however, a quantitative link between the observed dynamics and the surrounding chromatin structure could not yet be established in real-time. Although it is becoming increasingly clear that chromatin motion and long-range interactions are key to genome organization and gene regulation (*13*), tools to detect and to define bulk chromatin motion simultaneously at divergent spatio-temporal scales and high resolution are still missing.

Here we apply deep learning-based photo-activated localization microscopy (Deep-PALM) for temporally resolved super-resolution imaging of chromatin *in vivo*. Deep-PALM acquires a single resolved image in a few hundred milliseconds with a spatial resolution of ~ 60 nm. We observed elongated ~ 45 to 90 nm wide chromatin domain ‘blobs’. Employing a computational chromosome model, we inferred that blobs are highly dynamic entities, which dynamically assemble and disassemble. Consisting of chromatin in close physical and genomic proximity, our chromosome model indicates that blobs nevertheless adopt TAD-like interaction patterns when chromatin configurations are averaged over time. Using a combination of Deep-PALM and high-resolution dense motion reconstruction (*14*), we simultaneously analyzed both structural and dynamic properties of chromatin. Our analysis emphasizes the presence of spatio-temporal cross-correlations between chromatin structure and dynamics, extending several micrometers in space and tens of seconds in time. Furthermore, extraction and statistical mapping of multiple parameters from the dynamic behavior of chromatin blobs shows that chromatin density regulates local chromatin dynamics.

## Results

### Deep-PALM reveals dynamic chromatin remodeling in living cells

Super-resolution imaging of complex and compact macromolecules such as chromatin requires dense labeling of the chromatin fiber in order to resolve fine features. We employ Deep-STORM, a method which uses a deep convolutional neural network (CNN) to predict super-resolution images from stochastically blinking emitters (*15*) (Figure 1A; Materials and Methods). The CNN was trained to specific labeling densities for live-cell chromatin imaging using a photo-activated fluorophore (PATagRFP); we therefore refer to the method as Deep-PALM. We chose three labeling densities 4, 6 and 9 emitters per μm^2^ per frame in the ON-state to test, based on the comparison of simulated and experimental wide field images (Supplementary Figure 1A). The CNN trained with 9 emitters per μm^2^ performed significantly worse than the other CNNs and was thus excluded from further analysis (Supplementary Figure 1B; Materials and Methods). We applied Deep-PALM to reconstruct an image set of labeled histone protein (H2B-PATagRFP) in human bone osteosarcoma (U2OS) cells using the networks trained on 4 and 6 emitters per μm^2^ per frame (Materials and Methods). A varying number of predictions by the CNN of each individual frame of the input series were summed to reconstruct a temporal series of super resolved images (Supplementary Figure 1C). The predictions made by the CNN trained with 4 emitters per μm^2^ show large spaces devoid of signal intensity, especially at the nuclear periphery, making this CNN inadequate for live-cell super-resolution imaging of chromatin. While collecting photons from long acquisitions for super-resolution imaging is desirable in fixed cells, Deep-PALM is a live imaging approach. Summing over many individual predictions leads to considerable motion blur and thus loss in resolution. Quantitatively, the Nyquist criterion states that the image resolution 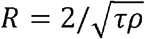 depends on *ρ*, the localization density per second and the time resolution τ (*16*). In contrast, motion blur strictly depends on the diffusion constant *D* of the underlying structure 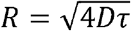. There is thus an optimum resolution due to the tradeoff between increased emitter sampling and the avoidance of motion blur, which was at a time resolution of 360 ms for our experiments (Figure 1B; Supplementary Figure 1D).

**Figure 1:**
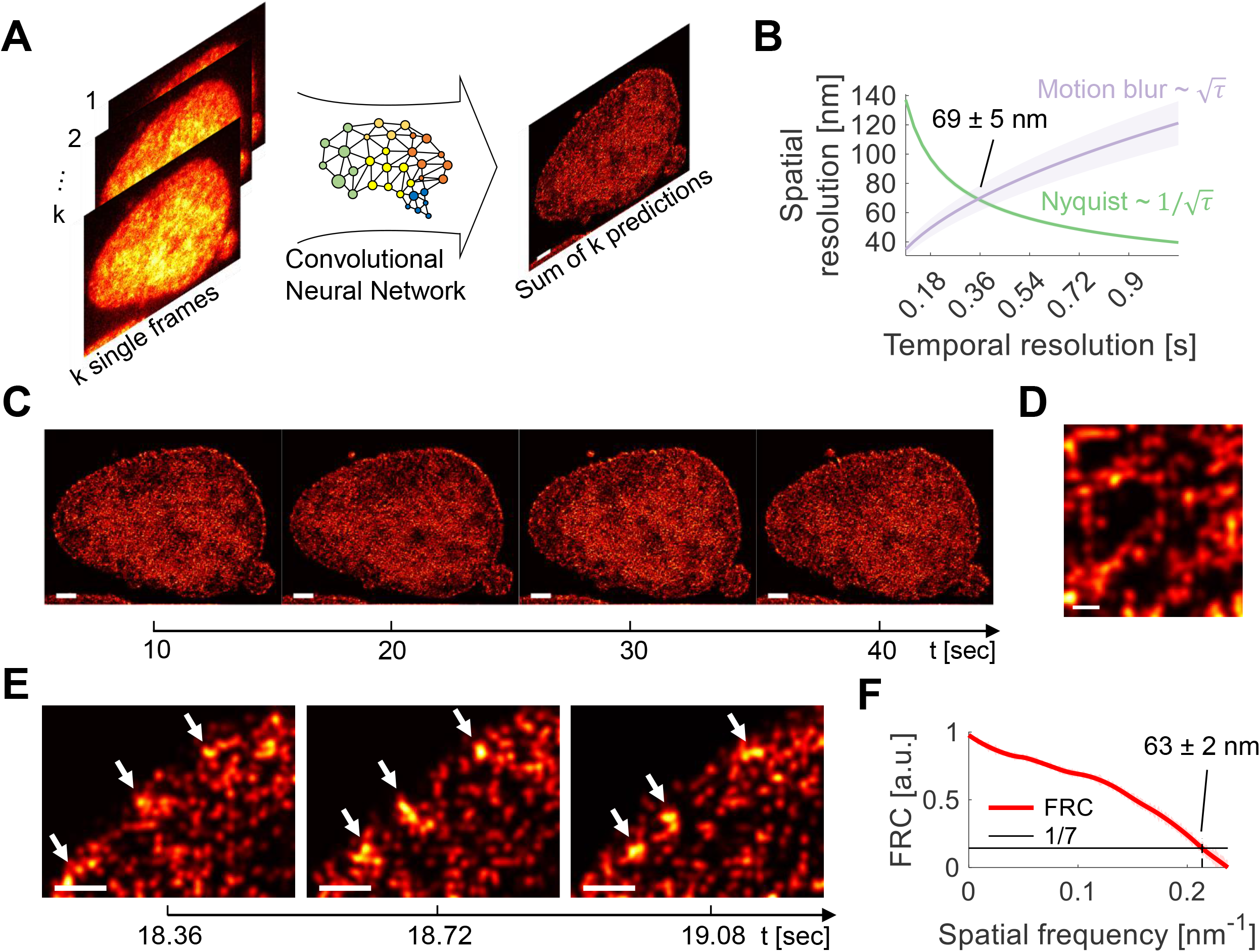
Temporally resolved super-resolution images of chromatin in U2OS nuclei. **A)** Widefield images of U2OS nuclei expressing H2B-PATagRFP are input to a trained convolutional neural network (CNN) and predictions from multiple input frames are summed to construct a super-resolved image of chromatin *in vivo.* **B)** The resolution tradeoff between the prolonged acquisition of emitter localizations (green line) and motion blur due to diffusion of the underlying diffusion processes (purple line). For our experimental data, the localization density per second is *ρ* = (2.4 ± 0.1) μm^−2^ s^−1^, the diffusion constant is *D* = (3.4 ± 0.8) · 10^−3^ μm^2^s^−1^ (see Supplementary Figure 8B) and the acquisition time per frame is *τ* = 30 ms. The spatial resolution assumes a minimum (69 ± 5 nm) at a time resolution of 360 ms. **C)** Super-resolution images of a single nucleus at time intervals of about 10 seconds. Scale bar is 2 μm. **D)** Magnification of segregated accumulations of H2B within a chromatin-rich region. Scale bar is 200 nm. **E)** Magnification of a stable, but dynamic structure (arrows) over three consecutive images. Scale bar is 500 nm. **F)** Fourier Ring Correlation (FRC) for super-resolved images resulting in a spatial resolution of 63 ± 2 nm. FRC was conducted based on 332 consecutive super resolved images from two cells.

Super-resolution imaging of H2B-PATagRFP in live cells at this temporal resolution shows a pronounced nuclear periphery while fluorescent signals in the interior vary in intensity (Figure 1C). This likely corresponds to chromatin-rich and chromatin-poor regions (*8*). These regions rearrange over time, reflecting the dynamic behavior of bulk chromatin. Chromatin-rich and chromatin-poor regions were visible not only at the scale of the whole nucleus, but also at the resolution of a few hundred nanometers (Figure 1D). Within chromatin-rich regions, the intensity distribution was not uniform but exhibited spatially segregated accumulations of labeled histones of variable shape and size, reminiscent of nucleosome clutches (*6*), nanodomains (*9*, *11*) or TADs (*17*). At the nuclear periphery, prominent structures arise. Certain chromatin structures could be observed for ~ 1 s, which underwent conformational changes during this period (Figure 1E). The spatial resolution at which structural elements can be observed (Materials and Methods) in time-resolved super-resolution data of chromatin was 63 ± 2 nm (Figure 1E), slightly more optimistic than the theoretical prediction (Figure 1B) (*18*).

We compared images of H2B reconstructed from 12 frames (super resolved images) by Deep-PALM in living cells to super-resolution images reconstructed by 8,000 frames of H2B in fixed cells (Supplementary Figure 2A, B). Overall, the contrast in the fixed sample appears higher and the nuclear periphery appears more prominent than in images from living cells. However, in accordance with previous super-resolution images of chromatin in fixed cells (*6*, *8*, *9*, *11*, *17*) and Deep-PALM images, we observe segregated accumulations of signal throughout the nucleus. Thus, Deep-PALM identifies spatially heterogeneous coverage of chromatin as previously reported (*6*, *8*, *9*, *11*, *17*). We further monitor chromatin temporally at nanometer scale in living cells.

### Chromatin appears in elongated nanometer-sized blobs with a non-random spatial distribution

To quantitatively assess the spatial distribution of H2B, we developed an image segmentation scheme (Materials and Methods, Supplementary Figure 3) which allowed us to segment spatially separated accumulations of H2B signal with high fidelity (Supplementary Note 1, Supplementary Figure 4, Supplementary Figure 5). Applying our segmentation scheme, ~ 10,000 separable elements, blob-like structures were observed for each super-resolved image (166 resolved images per movie; Figure 2A). The experimental resolution does not enable us to elucidate their origin and formation because tracking of blobs in three dimensions would be necessary to do so (see Discussion). We therefore turned to a transferable computational model introduced by Qi *et al.* (*19*), which is based on one-dimensional genomics and epigenomics data, including histone modification profiles and CTCF binding sites. To compare our data to the simulations, super-resolution images were generated from the modeled chromosomes. Within these images we could identify and characterize ‘chromatin blobs’ analogously to those derived from experimental data (Materials and Methods; Figure 2B).

**Figure 2:**
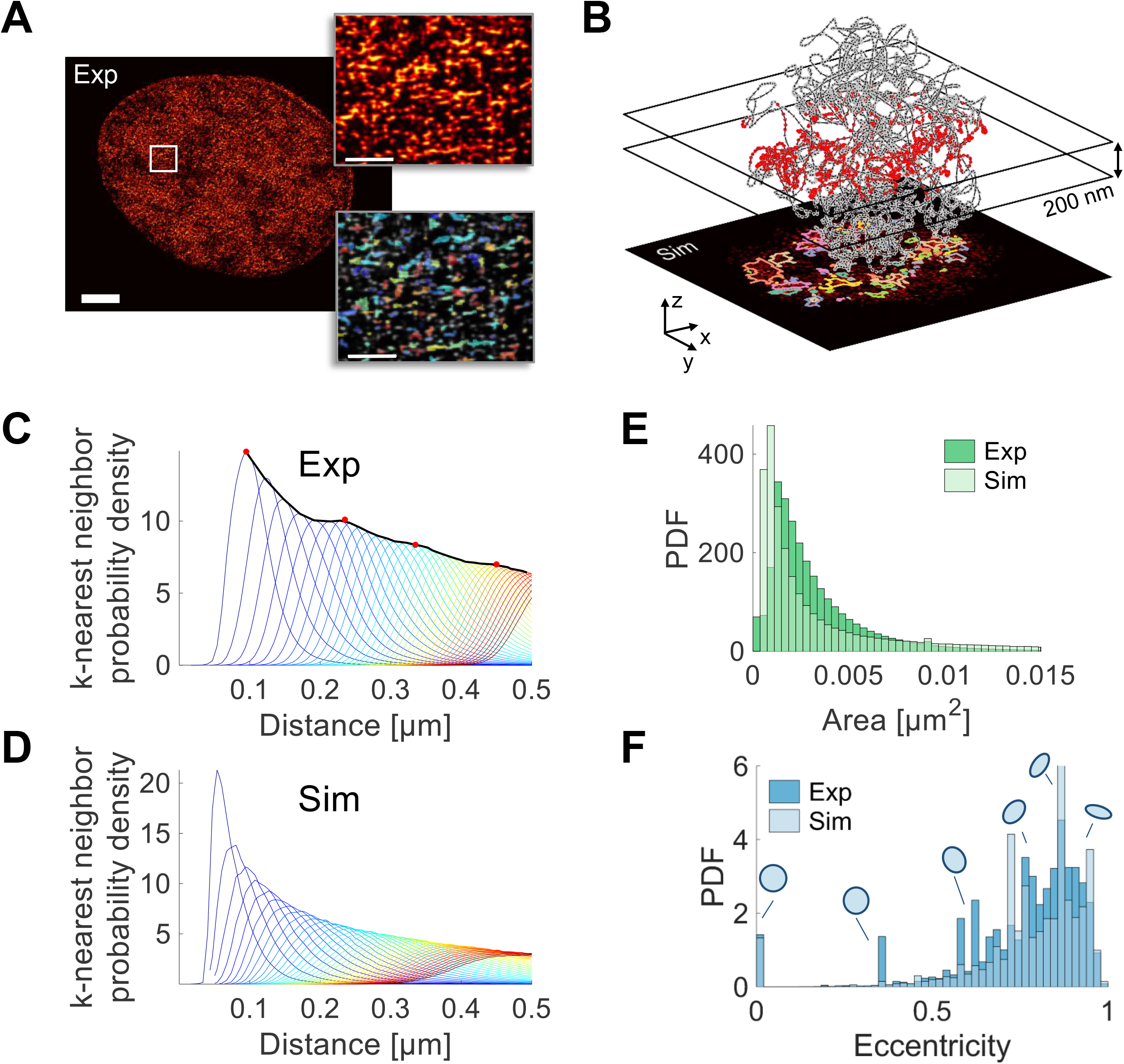
Chromatin blob identification and characterization of imaged and modeled chromatin. **A)** Super-resolved images show blobs of chromatin (left). These blobs are segmented (Materials and Methods, Supplementary Note 1) and individually labeled by random color (right). Magnifications of the boxed regions are shown. Scale bars: whole nucleus 2 μm, magnifications 200 nm. **B)** Generation of super-resolution images and blob identification and characterization for a 25 Mbp segment of chromosome 1 from GM12878 cells as simulated in Qi *et al.* (*19*). Beads (5 kb genomic length) of a simulated polymer configuration within a 200 nm thick slab are projected to the imaging plane, resembling experimental super-resolved images of live chromatin. Blobs are identified as on experimental data. **C)** From the centroid positions, the nearest-neighbor distance (NND) distributions are computed for up to 40 nearest neighbors (blue to red). The envelope of the k-NND distributions (black line) shows peaks at approximately 95 nm, 235 nm, 335 nm and 450 nm (red dots). **D)** k-NND distributions as in B) for simulated data. **E)** Area distribution of experimental and simulated blobs. The distribution is in both cases well described by a lognormal distribution with parameters (**3. 3** ± **2. 8**) · **1O**^−3^ ^2^ for experimental blobs and (**3. 1** ± **3. 2**) · **1O**^−3^ μ^2^ for simulated blobs (mean ± standard deviation). **F)** Eccentricity distribution for experimental and simulated chromatin blobs. Selected eccentricity values are illustrated by ellipses with the corresponding eccentricity. Eccentricity values range from 0, describing a circle, to 1, describing a line. Prominent peaks arise due to the discretization of chromatin blobs in pixels. The data is based on 332 consecutive super resolved images from two cells, in each of with ~ 10,000 blobs were identified.

For imaged (in living and fixed cells) and modeled chromatin, we first computed the k^th^ nearest neighbor distance (NND; centroid-to-centroid) distributions, taking into account the nearest 1^st^ to 40^th^ neighbors (Figure 2C and Supplementary Figure 2C, D, blue to red). Centroids of nearest neighbors are (95±30) nm (mean ± standard deviation) apart, consistent with previous and our own super-resolution images of chromatin in fixed cells (*9*) and slightly further than what was found for clutches of nucleosomes (*6*). The envelope of all NND distributions (Figure 2C, black line) shows several weak maxima at ~ 95 nm, 235 nm, 335 nm, and 450 nm, which roughly coincide with the peaks of the 1^st^, 7^th^, 14^th^ and 25^th^ nearest neighbors respectively (Figure 2C, red dots). In contrast, simulated data exhibit a prominent first nearest neighbor peak at slightly smaller distance and higher-order NND distribution decay quickly and appear washed out (Figure 2D). This hints towards greater levels of spatial organization of chromatin *in vivo*, which is not readily recapitulated in the employed state-of-the-art chromosome model.

Next, we were interested in the typical size of chromatin blobs. Their area distribution (Figure 2E) fit a log-normal distribution with parameters (3.3±2.8 · 10^−3^ μm^2^ (mean ± standard deviation), which is in line with the area distribution derived from fixed samples (Supplementary Figure 2E) and modeled chromosomes. Notably, blob areas vary considerably as indicated by the high standard deviation and the prominent tail of the area distribution towards large values. Following this we calculated the eccentricity of each blob to resolve their shape (Figure 2F; Supplementary Figure 2F). The eccentricity is a measure of the elongation of a region reflecting the ratio of the longest chord of the shape and the shortest chord perpendicular to it (Figure 2F, illustrated shapes at selected eccentricity values). The distribution of eccentricity values shows an accumulation of values close to 1, with a peak value of ~ 0.9, which shows that most blobs have an elongated, fiber-like shape and are not circular. In particular, the eccentricity value of 0.9 corresponds to a ratio between the short and long axis of the ellipse of ~ ½ (Materials and Methods), which results, considering the typical area of blobs in experimental and simulated data, in roughly 92 nm long and 46 nm wide blobs on average. A highly similar value was found in fixed cells (Supplementary Figure 2F). The length coincides with the value found for the typical nearest-neighbor distance (Figure 2C; (95±30 nm). However, due to the segregation of chromatin into blobs, their elongated shape and their random orientation (Figure 2A), the blobs cannot be closely packed throughout the nucleus. We find that chromatin has a spatially heterogeneous density, occupying 5 - 60% of the nuclear area (Supplementary Figure 6A, B), which is supported by a previous electron microscopy study (*20*).

Blob dimensions derived from live-cell super-resolution imaging using Deep-PALM are consistent with those found in fixed cells, thereby further validating our method, and in agreement with previously determined size ranges (*6*, *9*). A previously published chromosome model based on Hi-C data (and thus not tuned to display blob-like structures per se) also displays blobs with dimensions comparable to those found here, in living cells. Taken together, these data strongly suggest the existence of spatially segregated chromatin structures in the sub-100 nm range.

### Chromatin blobs identified by Deep-PALM are coherent with sub-TADs

The simulations offer to track each monomer (chromatin locus) unambiguously, which is currently not possible to do from experimental data. Since the simulations show blobs comparable to those found in experiment (Figure 2), simulations help to indicate possible mechanisms leading to the observation of chromatin blobs. For instance, due to the projection of the nuclear volume onto the imaging plane, the observed blobs could simply be overlays of distant, along the one-dimensional genome, non-interacting genomic loci. To examine this possibility, we analyzed the gap length between beads belonging to the same blob along the simulated chromosome. Beads constitute the monomers of the simulated chromosome and each bead represents roughly 5 kb (*19*).

The analysis showed that the blobs are mostly made of consecutive beads along the genome, thus implying an underlying domain-like structure, similar to TADs (Figure 3A). Using the affiliation of each bead to an intrinsic chromatin state of the model (Figure 3B), it became apparent that blobs along the simulated chromosome consisting mostly of active chromatin are significantly larger than those formed by inactive and repressive chromatin (Figure 3C). These findings are in line with experimental results (*10*) and results from the simulations directly (*19*), thereby validating the projection and segmentation process.

**Figure 3:**
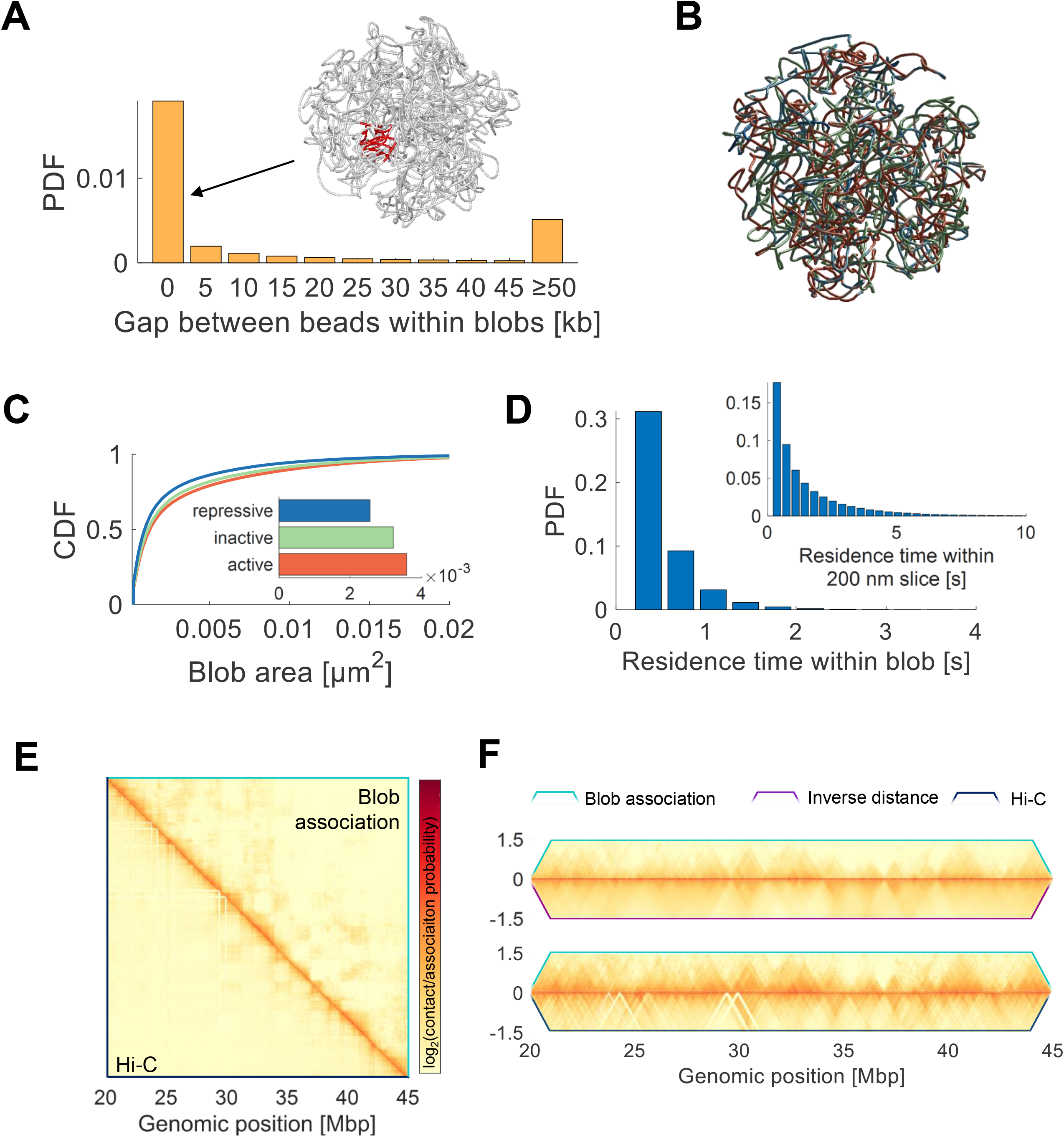
Chromatin blobs on modeled chromosomes consist of continuous loci along the genome and exhibit a TAD-like time-averaged conformation. **A)** Gap length between beads belonging to the same blob. An exemplary blob with small gap length is shown. The blob is mostly made of consecutive beads being in close spatial proximity. **B)** A representative polymer configuration is colored according to chromatin states (red: active, green: inactive, blue: repressive). **C)** The cumulative distribution function of clusters within active, inactive and repressive chromatin. Inset: Mean area of clusters within the three types of chromatin. The distributions are all significantly different from each other as determined by a two-sample Kolmogorov-Smirnov test (p < 10^−50^). **D)** Distribution of the continuous residence time of any monomer within a cluster (0.5 ± 0.3 s; mean ± standard deviation). Inset: Continuous residence time of any monomer within a slab of 200 nm thickness (1.5 ± 1.6 s; mean ± standard deviation). The blob association strength between any two beads is measured as the frequency at which any two beads are found in one blob. The association map is averaged over all simulated configurations (upper triangular matrix; from simulations) and experimental Hi-C counts are shown for the same chromosome segment (lower triangular matrix; from Rao *et al.* (*40*)). The association and Hi-C maps are strongly correlated (Pearson’s correlation coefficient PCC = 0.76) Close-up views around the diagonal of Hi-C-like matrices. The association strength is shown together with the inverse distance between beads (upper panel; PCC = 0.85) and with experimental Hi-C counts (lower panel; as in E)). The data is based on 20,000 polymer configurations.

Since chromatin is dynamic *in vivo* and in computer simulations, each bead can diffuse in and out of the imaging volume from frame to frame. We estimated that, on average, each bead spent approximately 1.5 s continuously within a slab of 200 nm thickness (Figure 3D). Furthermore, a bead is on average found only 0.55 ± 0.33 s continuously within a blob, which corresponds to 1 – 2 experimental super-resolved images (Figure 3D). These results suggest that chromatin blobs are highly dynamic entities, which usually form and dissemble within less than one second. We thus constructed a time-averaged association map for the modeled chromosomes, quantifying the frequency at which each locus is found with any other locus within one blob. The association map is comparable to interaction maps derived from Hi-C (Figure 3E). Strikingly, inter-loci association and Hi-C maps are strongly correlated, and the association map shows similar patterns as those identified as TADs in Hi-C maps, even for relatively distant genomic loci (> 1 Mbp). A similar TAD-like organization is also apparent when the average inverse distance between loci is considered (Figure 3F, upper panel), suggesting that blobs could be identified in super-resolved images due to the proximity of loci within blobs in physical space. The computational chromosome model indicates that chromatin blobs identified by Deep-PALM are mostly made of continuous regions along the genome and cannot be attributed to artifacts originating from the projection of the three-dimensional genome structure to the imaging plane. The simulations further indicate that the blobs associate and dissociate within less than one second, but loci within blobs are likely to belong to the same TAD. Their average genomic content is 75 kb, only a fraction of typical TAD lengths in mammalian cells (average size 880 kb) (*4*), suggesting that blobs likely correspond to sub-TADs or TAD nano-compartments (*17*).

### Quantitative chromatin dynamics at nanoscale resolution

To quantify the experimentally observed chromatin dynamics at the nanoscale, down to the size of one pixel (13.5 nm), we used a dense reconstruction of flow fields, Optical Flow (OF; Figure 4A, Materials and Methods), which was previously used to analyze images taken on confocal (*12*, *14*), and Structured Illumination Microscopes (*8*). We examined the suitability of OF for super-resolution based on single molecule localization images using simulations. We find that the accuracy of OF is slightly enhanced on super-resolved images, compared to conventional fluorescence microscopy images (Supplementary Note 2, Supplementary Figure 7A-C). Experimental super-resolution flow fields are illustrated on the basis of two subsequent images, between which the dynamics of structural features are apparent to the eye (Supplementary Figure 7D,E). On the nuclear periphery, connected regions spanning up to ~ 500 nm can be observed (Supplementary Figure 7D (i-ii), marked by arrows). These structures are stable for at least 360 ms but move from frame to frame. The flow field is shown on top of an overlay of the two super-resolved images and color-coded (Supplementary Figure 7D (iii), the intensity in frame 1 is shown in green, the intensity in frame 2 is shown in purple, co-localization of both is white). Displacement vectors closely follow the redistribution of intensity from frame to frame (roughly from green to purple). Similarly, structures within the nuclear interior (Supplementary Figure 7E) can be followed by eye, thus further validating and justifying the use of a dense motion reconstruction as a quantification tool of super-resolved chromatin motion.

**Figure 4:**
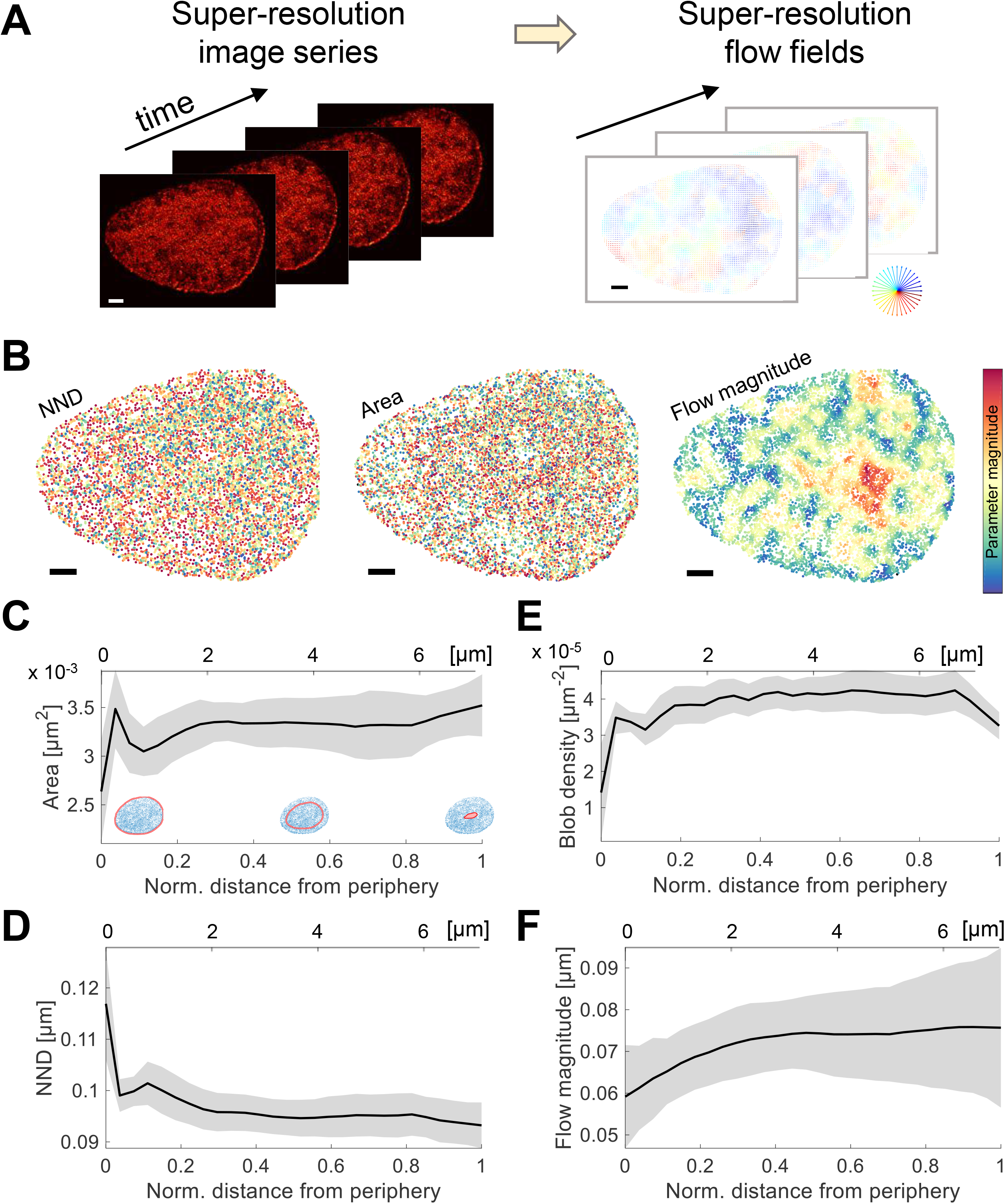
Structural and dynamics blob characteristics dependent on the proximity to the nuclear periphery. **A)** A time series of super-resolution images (left panel) is subject to Optical Flow (right panel). **B)** Blobs of a representative nucleus (see Supplementary Movie 1) are labeled by their NND (left), area (middle) and flow magnitude (right). Colors denote the corresponding parameter magnitude. **C)** The average blob area, **D)** NND, **E)** density and **F)** flow magnitude is shown versus the normalized distance from the nuclear periphery (lower x-axis; 0 is on the periphery and 1 is at the center of the nucleus) and versus the absolute distance (upper x-axis). Line and shaded area denote the mean ± standard error from 322 super resolved images of two cells.

Using Optical Flow fields, we linked the spatial appearance of chromatin to their dynamics. Effectively, the blobs were characterized with two structural parameters (NND and area) and their flow magnitude (Figure 4B). Supplementary Movie 1 shows the time evolution of those parameters for an exemplary nucleus. Blobs at the nuclear periphery showed a distinct behavior from those in the nuclear interior. In particular, the periphery exhibits a lower density of blobs, but those appear slightly larger and are less mobile than in the nuclear interior (Figure 4C-F), in line with previous findings using conventional microscopy (*14*). The peripheral blobs are reminiscent of dense and relatively immobile heterochromatin and lamina-associated domains (LADs) (*21*), which extend only up to 0.5 μm inside the nuclear interior. In contrast, blob dynamics increase gradually within 1 – 2 μm from the nuclear rim.

### Chromatin structure and dynamics are linked

To further elucidate the relationship between chromatin structure and dynamics, we analyzed the correlation between each pair of parameters in space and time. Therefore, we computed the auto- and cross-correlation of parameter maps with a given time lag across the entire nucleus (in space) (Figure 5A). In general, a positive correlation denotes a low-low or a high-high relationship (a variable de-/increases when another variable de-/increases) while analogously a negative correlation denotes a high-low relationship. The autocorrelation of NND maps (Figure 5A(i)) shows a positive correlation, thus regions exist spanning 2 - 4 μm, in which chromatin is either closely packed (low-low) or widely dispersed (high-high). Likewise, blobs of similar size tend to be in spatial proximity (Figure 5A(iii)). These regions are not stable over time but rearrange continuously, an observation bolstered by the fact that the autocorrelation diminishes with increasing time lag. The cross-correlation between NND and area (Figure 5A(ii)) shows a negative correlation for short time lags, suggesting that large blobs appear with a high local density while small ones are more isolated. Interestingly, the correlation becomes slightly positive for time lags ≥ 20 s, indicating that big blobs are present in regions which were sparsely populated before and small blobs tend to accumulate in previously densely populated regions. This is in line with dynamic reorganization and reshaping of chromatin domains on a global scale as observed in snapshots of the Deep-PALM image series (Figure 1A).

**Figure 5:**
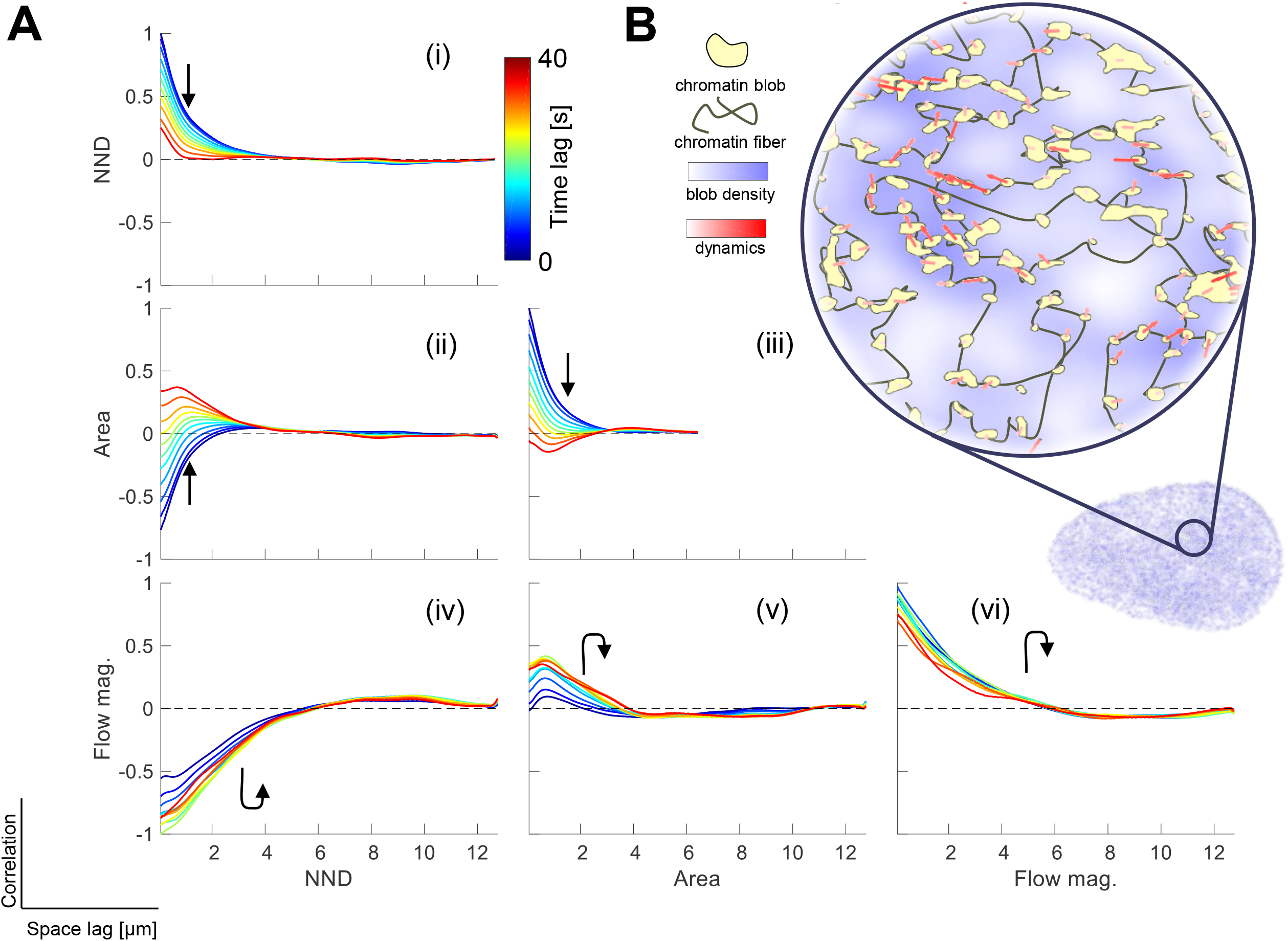
Spatio-temporal correlations between structural and dynamic parameters. **A)** The spatial auto- and cross-correlation between parameters were computed for different time lags. The graphs depict the correlation over space lag for each parameter pair and different colors denote the time lag (increasing from blue to red). **B)** Illustration of the instantaneous relationship between local chromatin density and dynamics. The blob density is shown in blue; the magnitude of chromatin dynamics is shown by red arrows. The consistent negative correlation between NND and flow magnitude is expressed by increased dynamics in regions of high local blob density. Data represents the average over two cells. The cells behave similarly such that error bars are omitted for the sake of clarity.

The flow magnitude is positively correlated for all time lags, while the correlation displays a slight increase for time lags ≤ 20 s (Figure 5A(vi)), which has also been observed previously (*8*, *12*, *22*). The spatial autocorrelation of dynamic and structural properties of chromatin are in stark contrast. While structural parameters are highly correlated at short, but not at long time scales, chromatin motion is still correlated at a time scale exceeding 30 s. At very short time scales (< 100 ms), stochastic fluctuations determine the local motion of the chromatin fiber, while coherent motion becomes apparent at longer times (*22*). However, there exists a strong cross-correlation between structural and dynamic parameters: the cross-correlation between the NND and flow magnitude shows striking negative correlation at all time lags (Figure 5A(iv)), strongly suggesting that sparsely distributed blobs appear less mobile than densely packed ones. The area seems to play a negligible role for short time lags, but there is a modest tendency that regions with large blobs tend to exhibit increased dynamics at later time points (≥ 10 s; Figure 5A(v)), likely due to the strong relationship between area and NND.

In general, parameter pairs involving chromatin dynamics exhibit an extended spatial auto-or cross-correlation (up to ~ 6 μm; the lower row of Figure 5A), compared to correlation curves including solely structural parameters (up to 3 - 4 μm). Furthermore, the cross-correlation of flow magnitude and NND does not considerably change for increasing time lag, suggesting that the coupling between those parameters is characterized by a surprisingly resilient memory, lasting for at least tens of seconds (*23*). Concomitantly, the spatial correlation of time-averaged NND maps and maps of the local diffusion constant of chromatin for the entire acquisition time enforce their negative correlation at the time scale of ~ 1 min (Supplementary Figure 8). Such resilient memory was also proposed by a computational study that observed that interphase nuclei behave like concentrated solutions of unentangled ring polymers (*24*). Our data support the view that chromatin is mostly unentangled since entanglement would influence the anomalous exponent of genomic loci in regions of varying chromatin density (*24*). However, our data do not reveal a correlation between the anomalous exponent and the time-averaged chromatin density (Supplementary Figure 8), in line with our previous results using conventional microscopy (*14*).

Overall, the spatial cross-correlation between chromatin structure and dynamics indicates that the NND between blobs and their mobility stand in a strong mutual, negative, relationship. This relationship, however, concerns chromatin density variations at the nanoscale, but not global spatial density variations such as in eu-or heterochromatin (*14*). These results support a model in which regions with high local chromatin density, i.e. larger blobs are more prevalent and are mobile, while small blobs are sparsely distributed and less mobile (Figure 5B). Blob density and dynamics in the long-time limit are to a surprisingly large extent influenced by preceding chromatin conformations.

### The local chromatin density is a key regulator of instantaneous chromatin dynamics

The spatial correlations above were only evaluated pairwise, while the behavior of every blob is likely determined by a multitude of factors in the complex energy landscape of chromatin (*19*, *22*). Here, we aim to take a wider range of available information into account in order to reveal the principle parameters, driving the observed chromatin structure and dynamics. Employing a microscopy-based approach, we have access to a total of six relevant structural, dynamic and global parameters, which potentially shape the chromatin landscape in space and time (Figure 6A). In addition to the parameters used above, we included the confinement level as a relative measure, allowing the quantification of transient confinement (Materials and Methods). We further included the bare signal intensity of super-resolved images and, as the only static parameter, the distance from the periphery since it was shown that dynamic and structural parameters show some dependence on this parameter (Figure 4). We then employed t-Distributed Stochastic Neighbor Embedding (*25*) (t-SNE), a state of the art dimensionality reduction technique, to map the six-dimensional chromatin ‘features’ (the six input parameters) into two dimensions. (Figure 6A, see Supplementary Note 3). The t-SNE algorithm projects data points such that neighbors in high-dimensional space likely stay neighbors in two-dimensional space (*25*). Visually apparent grouping of points (Figure 6B) implies that grouped points exhibit great similarity with respect to all input features and it is of interest to reveal which subset of the input features can explain the similarity among chromatin blobs best. It is likely that points appear grouped because their value of a certain input feature is considerably higher or lower than the corresponding value of other data points. We hence labeled points in t-SNE maps which are smaller than the first quartile point or larger than the third quartile point. Data points falling in either of the low/high partition of one input feature are colored accordingly for visualization (Figure 6D; blue/red points respectively). We then assigned a rank to each of the input features according to their nearest-neighbor fraction (n-n fraction): Since the t-SNE algorithm conserves nearest neighbors, we described the extent of grouping in t-SNE maps by the fraction of nearest neighbors which fall in either one of the subpopulations of low or high points (illustrated in Supplementary Figure 9). A high nearest neighbor fraction (n-n fraction; Figure 6C) therefore indicates that many points marked as low/high are indeed grouped by t-SNE and are therefore similar. The ranking (from low to high n-n fraction) reflects the potency of a given parameter to induce similar behavior between chromatin blobs with respect to all input features.

**Figure 6:**
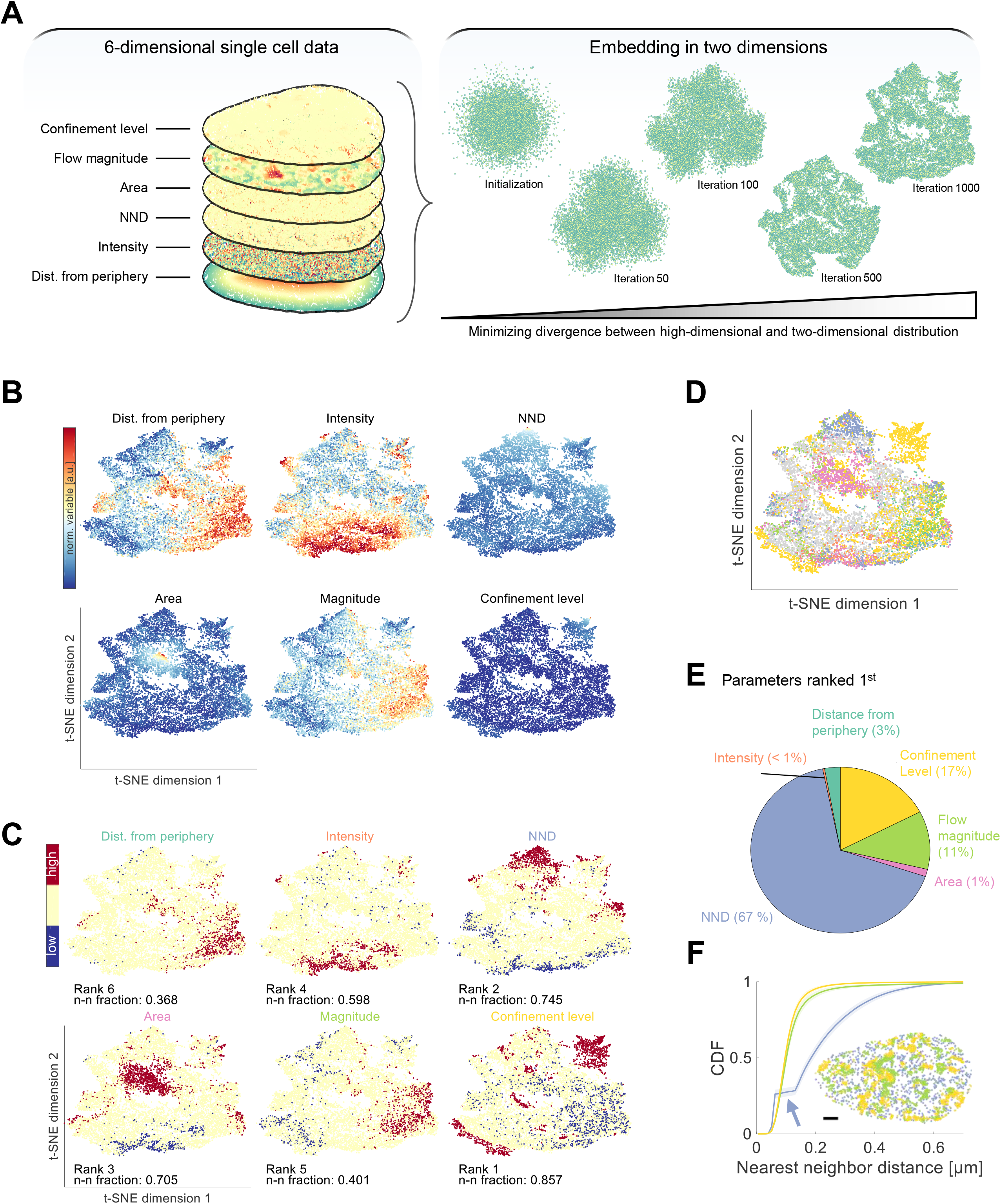
Chromatin feature extraction. **A)** The six-dimensional parameter space is input to the t-SNE algorithm and projected to two dimensions. **B)** The 2D embedding of an exemplary data set is shown and colored according to the magnitude of each input feature (blue to red, the parameter average is shown in beige) **C)** Points below the first (blue) and above the third (red) quartile points of the corresponding parameter are marked and the parameters are ranked according to the fraction of nearest neighbors which fall in one of the marked regions. **D)** Data points marked below the first or above the third quartile points are labeled according to the feature in which they were marked. Priority is given to the feature with the higher nearest-neighbor fraction if necessary. **E)** t-SNE analysis is carried out for each nucleus over the whole time series and it is counted how often a parameter ranked 1^st^. The results are visualized as a pie chart. The NND predominantly ranks 1^st^ in about 2/3 of all cases. **F)** Marked points in C-D) are mapped back onto the corresponding nuclei and the cumulative distribution function (CDF) over space is shown (mean ± standard error). Pie chart and CDF computations are based on 322 super resolved images from two cells.

The relative frequency at which each parameter ranked first provides an intuitive feeling for the most ‘influential’ parameters in the dataset (Figure 6E). The signal intensity plays a negligible role, suggesting that our data is free of potential artifacts related to the bare signal intensity. Furthermore, the blob area and the distance from the periphery likewise do not considerably shape chromatin blobs. In contrast, the NND between blobs was found to be the main factor inducing the observed characteristics in 67 % of all-time frames across all nuclei. The flow magnitude and confinement level together rank 1^st^ in 26 % of all cases (11 % and 17 %, respectively). These numbers suggest that the local chromatin density is a universal key regulator of instantaneous chromatin dynamics. Note that no temporal dependency is included in the t-SNE analysis and thus the feature extraction concerns only short-term (≤ 360 ms) relationships. The characteristics of roughly one-fourth of all blobs at each time point are mainly determined by similar dynamical features. Mapping chromatin blobs as marked in Figure 6C, *D* back to their respective positions inside the nucleus (Figure 6F) shows that blobs with low/high flow magnitude or confinement level markedly also grouped in physical space, which is highly reminiscent of coherent motion of chromatin (*12*). In contrast, blobs with extraordinary low or high NND were found interspersed throughout the nucleus, in line with spatial correlation analysis between structural and dynamic features (Figure 5). Our results point towards a large influence of the local chromatin density on the dynamics of chromatin at the scale of a few hundred nanometers and within a few hundred milliseconds. At longer time and length scales, however, previous results suggest that this relationship is lost (*14*).

## Discussion

With Deep-PALM we present temporally resolved super-resolution images of chromatin in living cells. Our technique identified chromatin nanodomains, named “blobs”, which mostly have an elongated shape, consistent with the curvilinear arrangement of chromatin as revealed by Structured Illumination Microscopy (*8*) with typical axes length of 45 – 90 nm. A previous study reported ~30 nm wide ‘clutches of nucleosomes’ in fixed mammalian cells using STORM nanoscopy (*6*), while the larger value obtained using Deep-PALM may be attributed to the motion blurring effect in live-cell imaging. However, histone acetylation and methylation marks were shown to form nanodomains of diameter 60 – 140 nm, respectively (*9*), which includes the computed dimensions for histone H2B using Deep-PALM.

To elucidate the origin of chromatin blobs, we turned to a simulated chromosome model, which displays chromatin blobs similar to our experimental data when seen in a super-resolution reconstruction. The simulations suggest that chromatin blobs consist of continuous genomic regions with an average length of 75 kb, assembling and disassembling dynamically within less than one second. Monomers within blobs display a distinct TAD-like association pattern in the long-time limit, suggesting that the identified blobs represent sub-TADs. Transient formation is consistent with recent findings that TADs are not stable structural elements, but exhibit extensive heterogeneity and dynamics (*3*, *5*). To experimentally probe the transient assembly of chromatin blobs, it would be interesting to track individual blobs over time. However, this is a non-trivial task. While the size (area/volume) or shape of blobs could be used to establish correspondences between blobs in subsequent frames, the framework needs to be flexible enough to allow for blob deformations since blobs likely arise stochastically and are not rigid bodies. Achieving an even shorter acquisition time per frame in the future could help minimize the influence of blob deformations and make tracking feasible. The second challenge is to distinguish between disassembly and out of focus diffusion of a blob. Three-dimensional imaging at sufficient spatial and temporal resolution will be helpful in the future to overcome this hurdle.

Using an Optical Flow approach to determine the blob dynamics instead, we found that structural and dynamic parameters exhibit extended spatial and temporal (cross-) correlations. Structural parameters such as the local chromatin density (expressed as the NND between blobs) and area lose their correlation after 3 – 4 μm and roughly 40 s in the spatial and temporal dimension, respectively. In contrast, chromatin mobility correlations extend over ~ 6 μm and persist during the whole acquisition period (≥ 40 s). Extensive spatio-temporal correlation of chromatin dynamics have been presented previously, both experimentally (*12*) and in simulations (*22*), but was not linked to the spatio-temporal behavior of the underlying chromatin structure until now. We found that the chromatin dynamics are closely linked to the instantaneous, but also to past local structural characterization of chromatin. In other words, the instantaneous local chromatin density influences chromatin dynamics in the future and vice versa. Based on these findings, we suggest that chromatin dynamics exhibit an extraordinary long memory. This strong temporal relationship might be established by the fact that stress propagation is affected by the folded chromosome organization (*26*). Fiber displacements cause structural reconfiguration, ultimately leading to a local amplification of chromatin motion in local high-density environments. This observation is also supported by the fact that increased nucleosome mobility grants chromatin accessibility even within regions of high nucleosome density (*27*).

Given the persistence at which correlations of chromatin structure and, foremost, dynamics occur in a spatio-temporal manner, we speculate that the interplay of chromatin structure and dynamics could involve a functional relationship (*28*): transcriptional activity is closely linked to chromatin accessibility and the epigenomic state (*29*). Because chromatin structure and dynamics are related, dynamics could also correlate with transcriptional activity (*14*, *30*, *31*). However, it is currently unknown if the structure-dynamics relationship revealed here is strictly mutual or if it may be causal. Simulations hint that chromatin dynamics follows from structure (*22*, *23*), this question will be exciting to answer experimentally and in the light of active chromatin remodelers in order to elucidate a potential functional relationship to transcription. Chromatin regions which are switched from inactive to actively transcribing, for instance, undergo a structural reorganization accompanied by epigenetic modifications (*32*). The mechanisms driving recruitment of enzymes inducing histone modifications such as histone acetyltransferases, deacetylases or methyltransferases is largely unknown, but often involves the association to proteins (*33*). Their accessibility to the chromatin fiber is inter alia determined by local dynamics (*27*). Such a structure-dynamics feedback loop would constitute a quick and flexible way to transiently alter gene expression patterns upon reaction to external stimuli or to co-regulate distant genes (*1*). Future work will study how structure-dynamics correlations differ in regions of different transcriptional activity and/or epigenomic states. Furthermore, to probe the interactions between key transcriptional machines such as RNA polymerases with the local chromatin structure and to record their (possibly collective) dynamics could shed light into the target search and binding mechanisms of RNA polymerases with respect to the local chromatin structure. Deep-PALM in combination with Optical Flow paves the way to answer these questions by enabling the analysis of time-resolved super-resolution images of chromatin in living cells.

## Materials and Methods

### Cell Culture

Human osteosarcoma U2OS expressing H2B-PATagRFP cells were a gift from Sébastien Huet (CNRS, UMR 6290, Institut Génétique et Développement de Rennes, Rennes, France); the histone H2B was cloned as described previously (*34*). U2OS cells were cultured in DMEM (with 4.5 g/l glucose) supplemented with 10% fetal bovine serum (FBS), 2 mM glutamine, 100 μg/ml penicillin, and 100 U/ml streptomycin in 5% CO_2_ at 37°C. Cells were plated 24 hours before imaging on 35 mm Petri dishes with a #1.5 coverslip like bottom (ibidi, Biovalley) with a density of 2×10^5^ cells/dish. Just before imaging, the growth medium was replaced by Leibovitz’s L-15 medium (Life Technologies) supplemented with 20% FBS, 2 mM glutamine, 100 μg/ml penicillin, and 100 U/ml streptomycin.

### PALM imaging in living cells

Imaging of H2B-PAtagRFP in living U2OS cells was carried out on a fully automated Nikon TI-E/B PALM (Nikon Instruments) microscope. The microscope is equipped with a full incubator enclosure with gas regulation to maintain a temperature of ~37°C for normal cell growth during live-cell imaging. Image sequences of 2,000 frames were recorded with an exposure time of 30◻ms per frame (33.3 frames/sec). For Deep-PALM imaging, a relatively low power (~50 W/cm2 at the sample) was applied for H2B-PATagRFP excitation at 561 nm and then combined with the 405 nm (~2 W/cm2 at the sample) to photo-activate the molecules between the states. Note that for Deep-PALM imaging, switched fluorophores are not required to stay as long in the dark state as for conventional PALM imaging. We used oblique illumination microscopy (*11*) combined with TIRF mode to illuminate a thin layer of 200 nm (axial resolution) across the nucleus. The reconstruction of super-resolved images improves the axial resolution only marginally (Supplementary Figure 1E, F). Laser beam powers were controlled by acoustic optic-modulators (AA Opto-Electronics). Both wavelengths were united into an oil immersion 1.49 NA TIRF objective (100x; Nikon). An oblique illumination was applied to acquire image series with high signal to noise ratio. The fluorescence emission signal was collected by using the same objective and spectrally filtered by a Quad-Band beam splitter (ZT405/488/561/647rpc-UF2, Chroma Technology) with Quad-Band emission filter (ZET405/488/561/647m-TRF, Chroma). The signal was recorded on an EMCCD camera (Andor iXon X3 DU-897, Andor Technologies) with a pixel size of 108 nm. For axial correction, Perfect Focus System was applied to correct for defocusing. NIS-Elements software was used for acquiring the images.

### PALM imaging and PALM data analysis in fixed cells

The same cell line (U2OS expressing H2B-PAtagRFP) as in live-cell imaging was used for conventional PALM imaging. Prior fixation, cells were washed with PBS (three times for 5 min each), then fixed with 4% paraformaldehyde (Sigma) diluted in PBS for 15 min at room temperature. A movie of 8,000 frames was acquired with an exposure time of 30 ms per frame (33.3 frames/sec). In comparison to Deep-PALM imaging , a relatively higher excitation laser of 561 nm (~60 W/cm^2^ at the sample) was applied to photo-bleach H2B-PATagRFP, then combined with the 405 nm (~2.5 W/cm^2^ at the sample) for photo-activating the molecules. We used the same oblique illumination microscopy combined with TIRF system as applied in living cells imaging.

PALM images from fixed cells were analyzed using ThunderSTORM (*35*). Super-resolution images were constructed by binning emitter localizations into 13.5 nm × 13.5 nm pixels and blurred by a Gaussian to match Deep-PALM images. The image segmentation was carried out as on images from living cells (see below).

### Deep-PALM analysis

The convolutional neural network (CNN) was trained using simulated data following Nehme *et al.* (*15*) for three labeling densities (4, 6 and 9 emitters per μm^2^ per frame). Raw imaging data were checked for drift as previously described (*12*). The detected drift in raw images is in the range < 10 nm and therefore negligible. The accuracy of the trained net was evaluated by constructing ground truth images from the simulated emitter positions. The Structural Similarity Index is computed to assess the similarity between reconstructed and ground truth images (*36*):

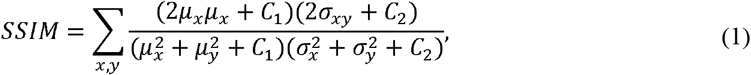

where *x*, and *y*, are windows of the predicted and ground truth images, respectively, *μ* and *σ* denote their local means and standard deviation, respectively and *σ*_*xy*_ their cross-variance. C_l_ = (0.01L)^2^ and C_2_ = (0.03L)^2^ are regularization constants, where *L* is the dynamic range of the input images. The second quantity to assess CNN accuracy is the Root Mean Square Error between the ground truth G and reconstructed image R:

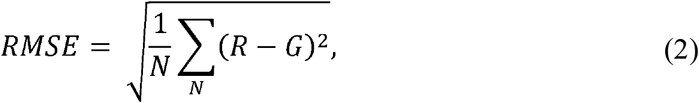

Where *N* is the number of pixels in the images. After training, sequences of all experimental images were compared to the trained network and predictions of single Deep-PALM images were summed to obtain a final super-resolved image. An up-sampling factor of 8 was used, resulting in an effective pixel size of 108 nm/8 = 13.5 nm. The image quality assessment in order to determine the optimal number of predictions to be summed, we use a blind/referenceless image spatial quality evaluator (BRISQUE) (*37*). For visualization, super-resolved images were convolved with a Gaussian kernel (*σ* = 1 pixel) and represented using a false RGB colormap. The parameters of the three trained networks are available at https://github.com/romanbarth/DeepPALM-trained-models

### Fourier Ring Correlation analysis

Fourier Ring Correlation (FRC) is an unbiased method to estimate the spatial resolution in microscopy images. We follow an approach similar to the one described in Nieuwenhuizen *et al.* (*38*). For localization-based super-resolution techniques, the set of localizations is divided into two statistically independent subsets and two images from these subsets are generated. The FRC is computed as the statistical correlation of the Fourier transforms of both sub-images over the perimeter of circles of constant frequency in the frequency domain. Deep-PALM, however, does not result in a list of localizations, but in predicted images directly. The set of 12 predictions from Deep-PALM were thus split into two statistically independent subsets and the method described in Nieuwenhuizen *et al.* (*38*) was applied.

### Chromatin blob identification

The super-resolved images displayed isolated regions of accumulated emitter density. To quantitatively assess the structural information implied by these accumulation of emitters in the focal plane, we developed a segmentation scheme which aims to identify individual blobs (Supplementary Figure 3). A marker-assisted watershed segmentation was adapted in order to accurately determine blob boundaries. For this purpose, we use the raw predictions from the deep convolutional neural network without convolution (Supplementary Figure 3A). The foreground in this image is marked by regional maxima and pixels with very high density (i.e. those with *I* > 0.99 *I*_max_, Supplementary Figure 3B). Since blobs are characterized by surrounding pixels of considerably less density, the Euclidian distance transform is computed on the binary foreground markers. Background pixels (i.e. those pixels not belonging to any blobs) are expected to lie far away from any blob center and thus, a good estimate for background markers are those pixels being furthest from any foreground pixel. We hence compute the watershed transform on the distance transform of foreground markers and the resulting watershed lines depict background pixels (Supplementary Figure 3C). Equipped with fore- and background markers (Supplementary Figure 3D), we apply a marker-controlled watershed transform on the gradient of the input image (Supplementary Figure 3E). The marker-controlled watershed imposes minima on marker pixels preventing the formation of watershed lines across marker pixels. Therefore, the marker-controlled watershed accurately detects boundaries as well as blobs which might not have been previously marked as foreground (Supplementary Figure 3F). Finally, spurious blobs whose median- or mean intensity is below 10% of the maximum intensity are discarded and each blob is assigned a unique label for further correspondence (Supplementary Figure 3G). The area and centroid position are computed for each identified blob for further analysis. This automated segmentation scheme performs considerably better than other state-of-the-art algorithms for image segmentation due to the reliable identification of fore- and background markers accompanied by the watershed transform (Supplementary Note 1).

### Chromatin blob properties

Centroid position, area, and eccentricity were computed. The eccentricity is computed by describing the blobs as an ellipse:

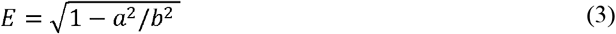

where *a* and *b* are the short and long axes of the ellipse, respectively.

### Chromatin blob identification from a computational chromatin model

We chose to employ a computational chromatin model, recently introduced by Qi *et al.* (*19*), in order to elucidate the origin of experimentally determined chromatin blobs. Each bead of the model covers a sequence length of 5 kb and is assigned one of 15 chromatin states to distinguish promoters, enhancers, quiescent chromatin, etc. Starting from the simulated polymer configurations, we consider monomers within a 200 nm thick slab through the center of the simulated chromosome. In order to generate super-resolved images as those from Deep-PALM analysis, fluorescence intensity is ascribed to each monomer. Monomer positions are subsequently discretized on a grid with 13.5 nm spacing and convolved with a narrow point-spread function, which results in images closely resembling experimental Deep-PALM images of chromatin. Chromatin blobs were then be identified and characterized as on experimental data (Figure 2A, B). Mapping back the association of each bead to a blob (if any) allows us to analyze principles of blob formation and maintenance using the distance and the association strength between each pair of monomers, averaged over all 20,000 simulated polymer configurations.

### Radial distribution function

The radial distribution function *g*(*r*) (RDF, also pair correlation function) is calculated (in two dimensions) by counting the number of blobs in an annulus of radius *r* and thickness *dr*. The result is normalized by the bulk density *ρ* = *n/A*, with the total number of blobs *n* and *A* the area of the nucleus, and the area of the annulus, 2*πr dr*:

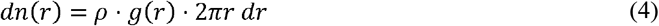

### Quantification of chromatin dynamics

Super-resolved images of chromatin showed spatially distributed blobs of varying size, but the resolved structure is too dense for state-of-the-art single particle tracking methods to track. Furthermore, are highly dynamic structures, assembling and dissembling within 1 – 2 super-resolved frames (Figure 3D), which makes a Single Particle Tracking approach unsuitable. Instead, we used a method for dynamics reconstruction of bulk macromolecules with dense labeling, Optical Flow. Optical Flow builds upon the computation of flow fields between two successive frames of an image series. The integration of these flow fields from super-resolution images results in trajectories displaying the local motion of bulk chromatin with temporal and high spatial resolution. Further, the trajectories are classified into various diffusion models and parameters describing the underlying motion are computed (*14*). Here, we use the effective diffusion coefficient *D* (in units of *m*^2^/*S*^α^), which reflects the magnitude of displacements between successive frames (the velocity of particles or monomers in the continuous limit) and the anomalous exponent *α* (*14*). The anomalous exponent reflects if the diffusion is free (*α* = 1, e.g. for non-interacting particles in solution), directed (*α* > 1, e.g. as the result from active processes) or hindered (*α* <1, e.g. due to obstacles or an effective back-driving force). Furthermore, we compute the length of constraint *L*_*c*_ which is defined as the standard deviation of the trajectory positions with respect to its time-averaged position. Denoting ***R***(*t*;***R***_0_) the trajectory at time *t* originating from ***R***_**0**_, the expression reads *L*_*c*_ = *var*(**R**(t;**R**_**0**_))^½^ where *var* denotes the variance. The length of constraint is a measure of the length scale explored of the monomer during the observation period. A complementary measure is the confinement level (*39*), which computes the inverse of the variance of displacements within a sliding window of length ω: *C* ∝ω/var(**R**(t;**R**_**0**_)), where the sliding window length is set to 4 frames (1.44 s). Larger values of *C* denote a more confined state than small ones.

### Spatial correlation for temporally varying parameters

The nearest-neighbor distance and the area, as well as the flow magnitude, were calculated and assigned to the blobs’ centroid position. In order to calculate the spatial correlation between parameters, the parameters were interpolated from the scattered centroid positions onto a regular grid spanning the entire nucleus. Because not every pixel in the original super-resolved images is assigned a parameter value, we chose an effective grid spacing of 5 pixels (67.5 nm) for the interpolated parameter maps. After interpolation, the spatial correlation was computed between parameter pairs: Let ***r*** = (*x,y*)^T^ denote a position on a regular two-dimensional grid and *f*(**r,t**) and *g*(**r**,*t*) two scalar fields with mean zero and variance one, at time *t* on that grid. The time series of parameter fields consist of *N* time points. The spatial cross-correlation between the fields *f* and *g*, which lie a lag time *τ* apart, is then calculated as

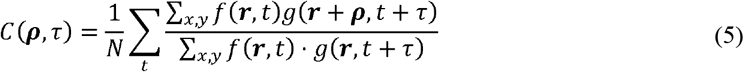

where the space lag *ρ* is a two-dimensional vector ***ρ*** = (Δ*x*,Δ*y*)^T^. The sums in the numerator and denominator are taken over the spatial dimensions, the first sum is taken over time. The average is thus taken over all time points which are compliant with time lag *τ*. Subsequently, the radial average in space is taken over the correlation, thus effectively calculating the spatial correlation *C*(*ρ,τ*) over the space lag 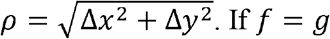 the spatial autocorrelation is computed.

### Spatial correlation for static parameters

We denote as global parameters those which reflect the structural and dynamic behavior of chromatin spatially resolved in a time-averaged manner. Examples involve the diffusion constant, the anomalous exponent, the length of constraint, but also time-averaged nearest-neighbor distance maps, etc. (Supplementary Figure 8). Those parameters are useful to determine time-universal characteristics. The spatial correlation between those parameters is equivalent to the expression given for temporally varying parameters when the temporal dimension is omitted, effectively resulting in a correlation curve *C*(*ρ*).

### t-Distributed Stochastic Neighbor Embedding (t-SNE)

The distance from the periphery, intensity, their nearest-neighbor distance, area, flow magnitude and confinement level of each identified blob form the six-dimensional input feature space for t-SNE analysis. The parameters for each blob (n = 3,260,232, divided into subsets of approximately 10,000) were z-transformed prior to the t-SNE analysis. The t-SNE analysis was performed using MATLAB and the Statistics and Machine Learning Toolbox (Release 2017b, The MathWorks, Inc., Natick, Massachusetts, United States) with the Barnes-Hut approximation. The algorithm was tested using different distance metrics and perplexity values and showed robust results within the examined ranges (Supplementary Note 3, Supplementary Figure 10).

### H2: Supplementary Materials

Note S1. Performance of the segmentation scheme employed for this study in comparison to other state-of-the-art algorithms for general purpose and comparison of blob segmentation on experimental images and on images containing randomly distributed emitters.

Note S2. Suitability of Optical Flow for super-resolution images of chromatin.

Note S3. t-SNE and its robustness with respect to distance metrics and perplexity values.

Figure S1. CNN training and time-resolution determination.

Figure S2. Comparison of Deep-PALM to super-resolution images of H2B in fixed U2OS cells.

Figure S3. Chromatin blob identification pipeline.

Figure S4. Performance of segmentation algorithms on super-resolved images of chromatin *in vivo*.

Figure S5. Segmentation on images of randomly distributed emitters.

Figure S6. Chromatin area fraction.

Figure S7. Performance of Optical Flow on conventional and super-resolved images and examples of super-resolution chromatin flow fields.

Figure S8. Global spatial correlation of structural and dynamic parameters.

Figure S9. Clustering illustration of points within a subset based on nearest-neighbors in t-SNE maps.

Figure S10. t-SNE for different distance metrics and perplexity values.

Movie S1. Time series of super-resolved chromatin structure and dynamics.

## Supporting information

yes

## Acknowledgments

**General**: We acknowledge support from the Pôle Scientifique de Modélisation Numérique, ENS de Lyon for providing computational resources. We thank Bin Zhang (Massachusetts Institute of Technology) for providing data of simulated chromosomes, Silvia Kocanova (LBME, CBI-CNRS; University of Toulouse) for providing PALM videos for fixed cells. We thank Hazen Babcock (Harvard University), Andrew Seeber (Harvard University) and Mikhail Tamm (Moscow State University) for valuable feedback on the manuscript. This publication is based upon work from COST Action CA18127, supported by COST (European Cooperation in Science and Technology).

## Author contributions

H.A.S. designed and supervised the project; R.B. designed the data analysis and wrote the code, H.A.S carried out experimental work; R.B. carried out the data analysis; H.A.S. and R. B interpreted results: H.A.S, R.B. and K.B. wrote the manuscript.

## Data and materials availability

All data needed to evaluate the conclusions in the paper are present in the paper and/or the Supplementary Materials. Additional data related to this paper may be requested from the authors.

