## Supplementary material for "Coupling chromatin structure and dynamics by live super-resolution imaging": yes

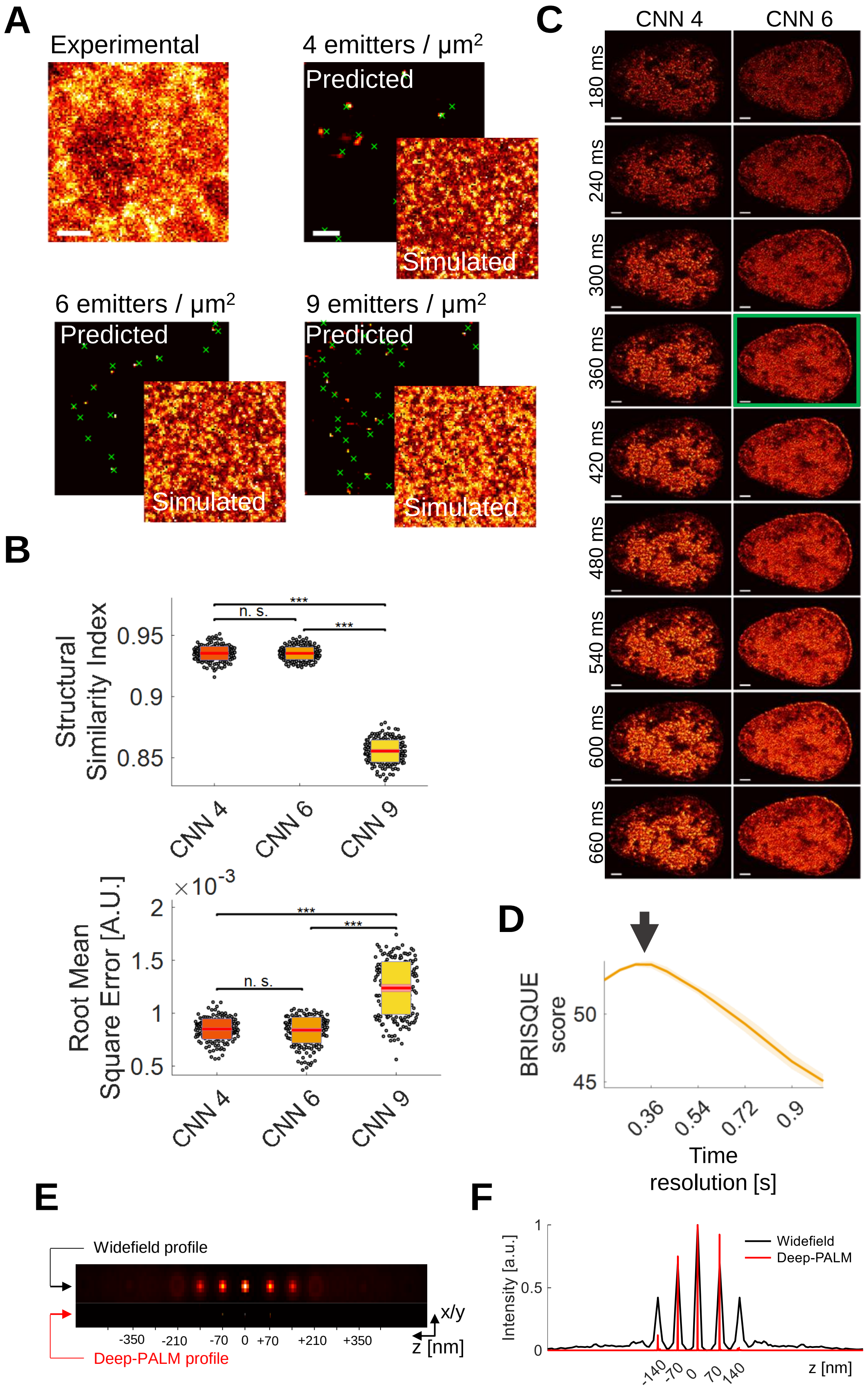
A
Experimental
4 emitters / μm2
Predicted
Simulated
6 emitters / μm2
Predicted
Simulated
9 emitters / μm2
Predicted
Simulated
B
Structural Similarity Index
n. s.
***
***
0.95
0.9
0.85
CNN 4
CNN 6
CNN 9
Root Mean Square Error [A.U.]
×10-3
2
1.5
1
0.5
n. s.
***
***
CNN 4
CNN 6
CNN 9
C
CNN 4
CNN 6
180 ms
240 ms
300 ms
360 ms
420 ms
480 ms
540 ms
600 ms
660 ms
D
BRISQUE score
50
45
0.36
0.54
0.72
0.9
Time resolution [s]
E
Widefield profile
Deep-PALM profile
-350
-210
-70
0
+70
+210
+350
x/y
z [nm]
F
Intensity [a.u.]
1
0.5
0
Widefield
Deep-PALM
-140
-70
0
70
140
z [nm]

**Supplementary Figure 1: CNN training and time-resolution determination. A)** Experimental and simulated widefield images with varying labeling density. Predictions by the trained CNNs are shown as false-color images and the ground truth emitter positions are overlaid as green crosses. Scale bar for experimental widefield image is 2 µm, simulated images are the same size. Scale bar on predictions are 200 nm. **B)** The accuracy of the trained CNNs was evaluated using the Structural Similarity Index and the Root Mean Square Error (Materials and Methods). The CNN trained with 9 emitters per $\mu m^2$ (CNN 9) performs significantly worse than the other two CNNs. Statistical significance assessed by a two-sample t-test (*** $p < 0.001$). **C)** Predictions from single acquired images are summed over different times from 360 ms to 1020 ms. A subset of reconstructions up to a time resolution of 660 ms are shown for CNN 4 and CNN 6. Scale bar is 2 µm. **D)** The structural image quality as quantified by BRISQUE (*1*) for CNN 6 at varying temporal resolution. The optimal (maximum) value is found at a time resolution of 360 ms. Data is shown as mean ± standard error from two cells. **E)** A point emitter is placed at different z distances from the focus (upper graph). The reconstruction of this emitter by the chosen CNN (lower graph) shows that, while the x-y resolution is greatly enhanced, the z-resolution is only marginally affected. **F)** Profiles across the widefield and reconstructed images of E) as indicated by arrows.

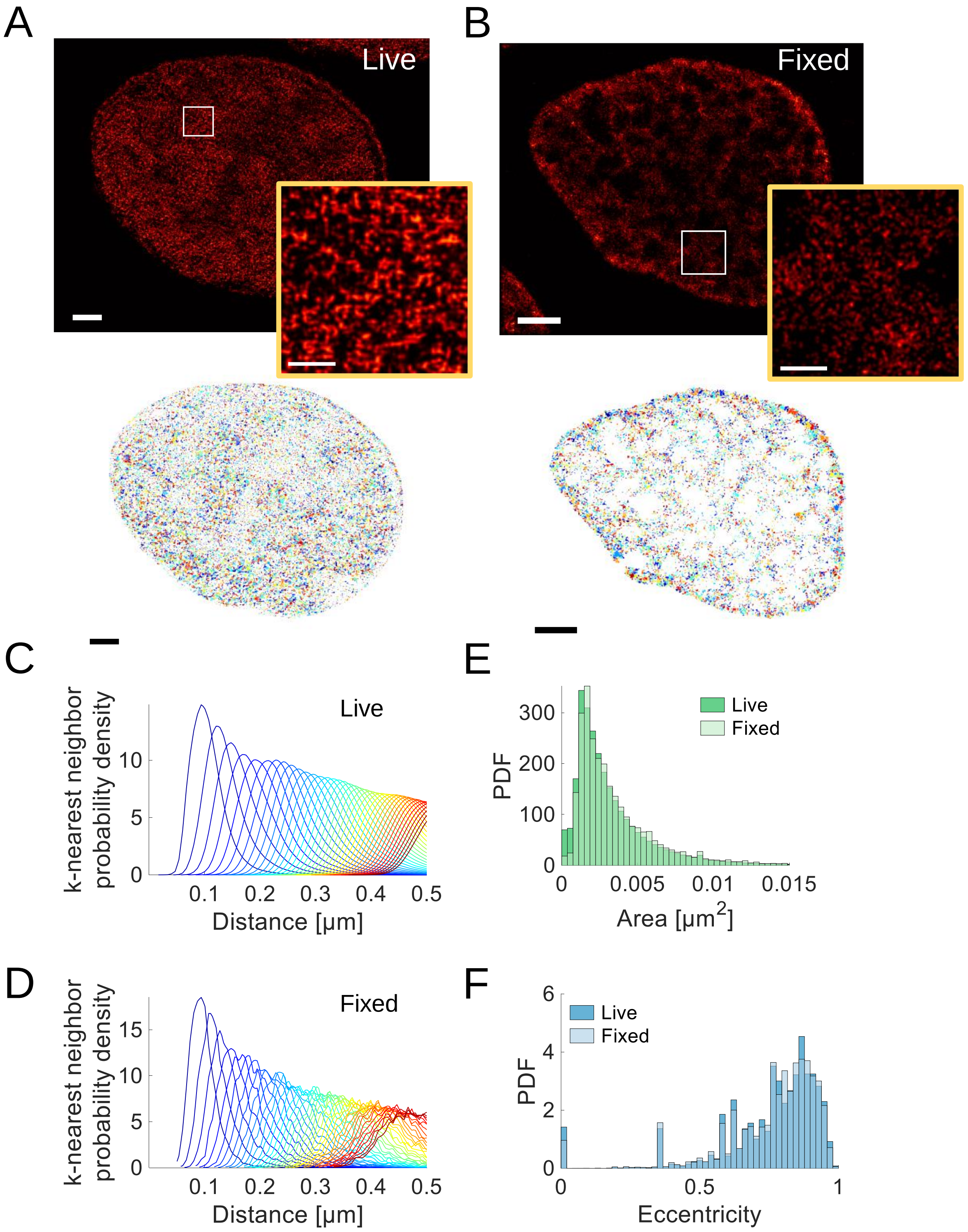


**Supplementary Figure 2: Comparison of Deep-PALM to "conventional" PALM images of H2B in fixed U2OS cells. A)** An exemplary image reconstructed by Deep-PALM with time resolution of 360 ms (12 frames) and the corresponding segmentation into chromatin blobs (lower panel; see Figure 2 and Supplementary Figure 3). **B)** An exemplary 'conventional' PALM image from 8000 frames of H2B in a fixed cell and the corresponding segmentation. The nuclear periphery is more pronounced in the fixed cell image, while the chromatin blobs are seen in both cases with comparable frequency (~ 10,000 per nucleus). Scale bars are 2 µm in original and 500 nm in magnified views. **C)** The nearest-neighbor distance distribution for the first 40 neighbors are shown from blue to red. Data as shown in Figure 2 for the live-cell imaging. Nearest neighbors are (95 ± 30) nm apart. **D)** The nearest neighbor distributions as in C) for the blobs detected in fixed cells (mean ± standard deviation: (88 ± 26) nm)). **E)** The area distribution for live and fixed cell imaging. The live-cell distribution is also shown in Figure 2. An average area of $(3.3 \pm 2.8) \cdot 10^{-3}$ $\mu m^2$ was found in living and $(3.5 \pm 2.5) \cdot 10^{-3}$ $\mu m^2$. **F)** The eccentricity distribution for blobs in living and fixed cells. The live-cell distribution is also shown in Figure 2. An average eccentricity of 0.75 ± 0.19 was found in living and 0.76 ± 0.18 in fixed cells. Appearance, NND, area and eccentricity distributions derived from either living or fixed samples correspond, thus validating the Deep-PALM live cell imaging approach. Data are shown for two living and two fixed cells.

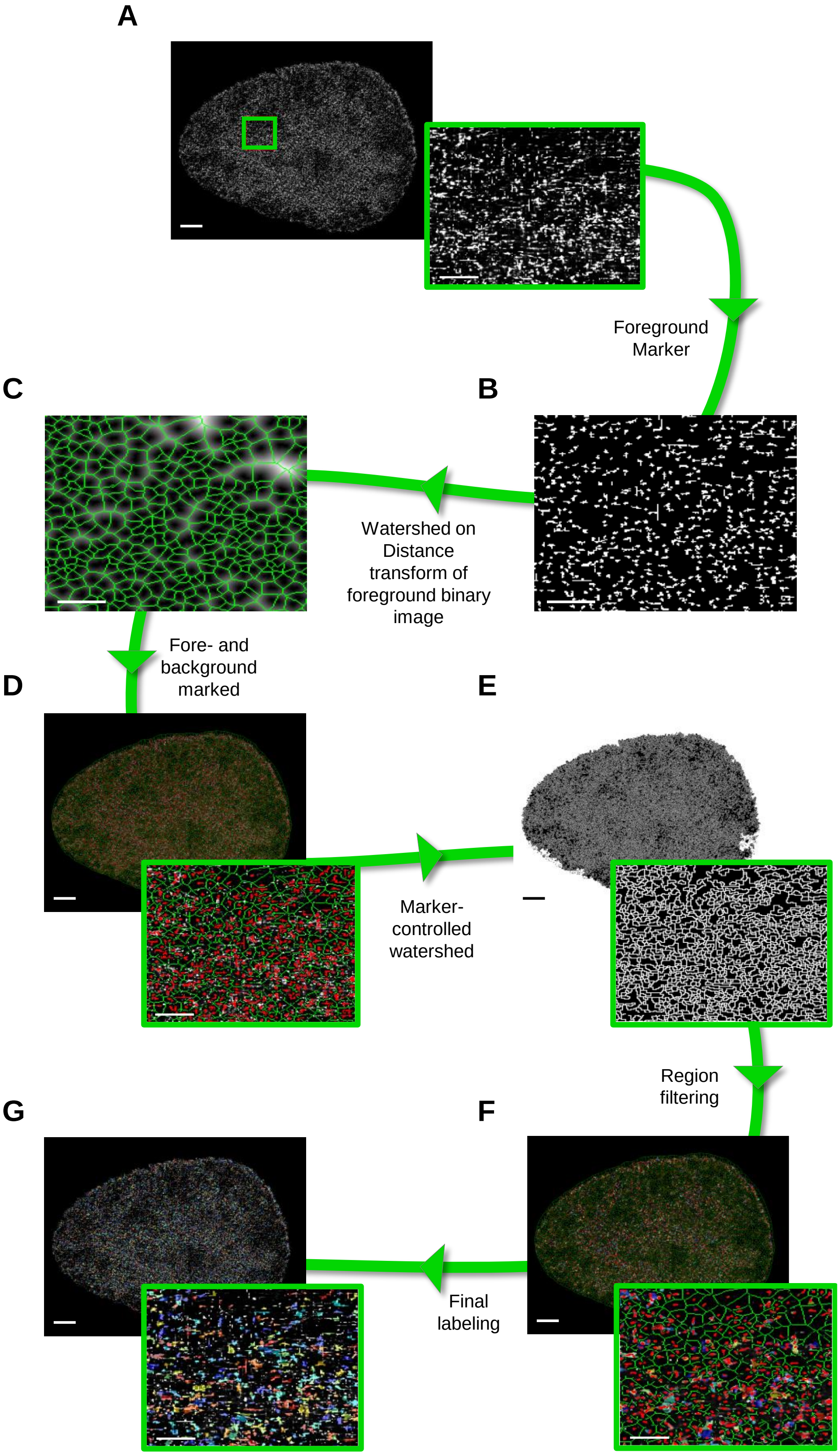
A
Foreground
Marker
C
B
Watershed on
Distance
transform of
foreground binary
image
Fore- and
background
marked
D
E
Marker-
controlled
watershed
Region
filtering
G
F
Final
labeling

**Supplementary Figure 3: Chromatin blob identification pipeline.** **A)** A grayscale image as output from the Deep-PALM algorithm is to be segmented. Magnified views correspond to the red rectangle. **B)** Foreground pixels are marked by finding the regional maxima. Additionally, very bright pixels (i.e. those with $\boldsymbol{I > 0.99 \cdot I_{max}}$) are marked as foreground. Only the magnification is shown. **C)** The watershed algorithm (without markers) is applied to the distance transform of the foreground mask. The resulting watershed lines represent lines in between pixels marked as foreground and therefore represent pixels of (local) low intensity, i.e. background markers. The background mask is corrected for pixels which nevertheless exert high intensity (i.e. those with $\boldsymbol{I > 0.8 \cdot I_{max}}$). Only the magnification is shown. **D)** Input image with the foreground (red) and background (green) markers superimposed. **E)** The watershed algorithm is applied to the gradient of the input image with minima imposed on pixels belonging to fore- or background. **F)** The regions defined by watershed lines (the basins) are post-processed in order to keep only those whose mean and median intensity exceed a threshold of $\boldsymbol{0.1 \cdot I_{max}}$. **G)** Resulting segmentation of the input image. Scale bars correspond to 2 µm and 0.5 µm in the full and magnified regions respectively.

### SUPPLEMENTARY NOTE 1

#### Segmentation performance

The marker-assisted watershed segmentation pipeline developed to segment chromatin blobs in super-resolved images is tested against other state-of-the-art segmentation algorithms in order to validate its performance. We evaluated our approach against three fully automated improvements of the widely spread watershed algorithm: the enhanced waterfall algorithm, the 'standard' algorithm, and the P-algorithm (*2*). To this end, we simulated images which closely resemble experimental super-resolved images of histone-labeled chromatin. We generated random shapes by thresholding randomly generated 1/f noise and gave intensity to the shapes according to experimental images. An exemplary simulated and experimental image is shown in Supplementary Figure 4A, B. The segmentation result of an exemplary simulated image is shown in Supplementary Figure 4D, I. We used the Rand index as a measure of the similarity of two segmentation algorithms. In particular, the Rand index is computed between the ground truth and the segmentation result of each algorithm as an estimate of the likelihood of a correctly classified element. An image segmentation problem can be formulated as a partitioning of the image pixels into several subsets. Let $P$ denote the set of $n$ pixels to be segmented, $G = \{G_1, \dots, G_g\}$ denotes the ground truth segmentation of $n$ pixels into $g$ subsets and $R = \{R_1, \dots, R_r\}$ denotes the resulting segmentation into $r$ subsets from an algorithm to test. The number of agreements between $G$ and $R$ consists of (i) the true positives (TP), i.e. the pairs of elements in $P$ that are in the same subset in $G$ and $R$ and (ii) the true negatives (TN), i.e. the pairs of elements in $P$ that are not in the same subset in $G$ and not in $R$. Likewise, the number of disagreements consists of (iii) the false negatives (FN), i.e. elements in $P$ that are in the same subset in $G$ but not in $R$ and (iv) the false positives (FP), i.e. elements in $P$ that are not in the same subset in $G$ but that are in $R$. Taken together, the Rand index expresses the number of agreements over the total number of pairs, i.e. agreements and disagreements (*3*):

$$\text{Rand Index} = \frac{TP + TN}{TP + TN + FN + FP} = \frac{TP + TN}{\binom{n}{2}} \tag{1}$$

The Rand index ranges from 0 to 1 and gives the fraction of matching pairs of pixels in the ground truth and computed segmentation. Our custom marker-controlled watershed algorithm performs significantly better than other tested algorithms, with a Rand index of about 75%.

To further validate the segmentation algorithm, the blob segmentation on experimental images of chromatin (Supplementary Figure 5A) was compared to images, in which emitters were randomly distributed (Supplementary Figure 5B). On random images, the blob density was ~ 19-fold reduced compared to segmentation on experimental images (Supplementary Figure 5C) and the blob area was about one order of magnitude smaller (Supplementary Figure 5D). These results show that blobs were identified due to the appearance of chromatin as blobs and not due to apparent grouping of randomly distributed emitters.

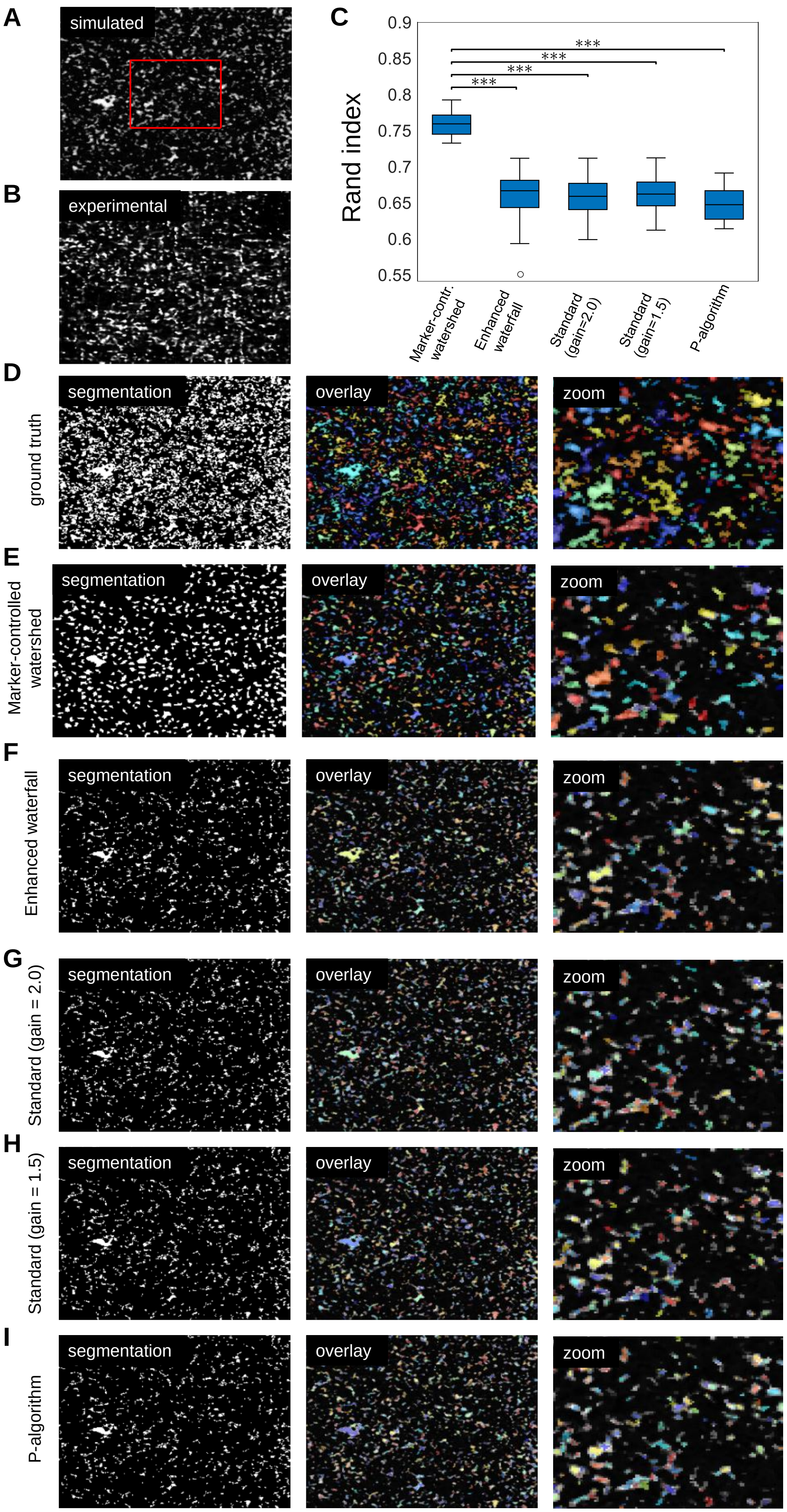
A
simulated
B
experimental
C
Rand index
0.9
0.85
0.8
0.75
0.7
0.65
0.6
0.55
***
***
***
***
Marker-contr. watershed
Enhanced waterfall
Standard (gain=2.0)
Standard (gain=1.5)
P-algorithm
D
ground truth
segmentation
overlay
zoom
E
Marker-controlled watershed
segmentation
overlay
zoom
F
Enhanced waterfall
segmentation
overlay
zoom
G
Standard (gain = 2.0)
segmentation
overlay
zoom
H
Standard (gain = 1.5)
segmentation
overlay
zoom
I
P-algorithm
segmentation
overlay
zoom

**Supplementary Figure 4: Performance of segmentation algorithms on super-resolved images of chromatin *in vivo*. A)** Exemplary simulated image for which the ground truth segmentation is known, used to compare different segmentation algorithms. The section in the red triangle is magnified in D-I). **B)** Exemplary experimental image to be compared to the simulated image. **C)** The Rand index reveals that our custom maker-controlled watershed algorithm significantly outperforms all other tested algorithms. Statistical significance assessed from 20 independent segmentation runs by a two-sample t-test (*** $p < 0.001$). **D)** Binary image (right) displaying the ground truth segmentation of the simulated image. Different segments are randomly colored for clarity and the magnified region marked in A) is shown. **E-I)** Exemplary segmentation results from the tested algorithms.

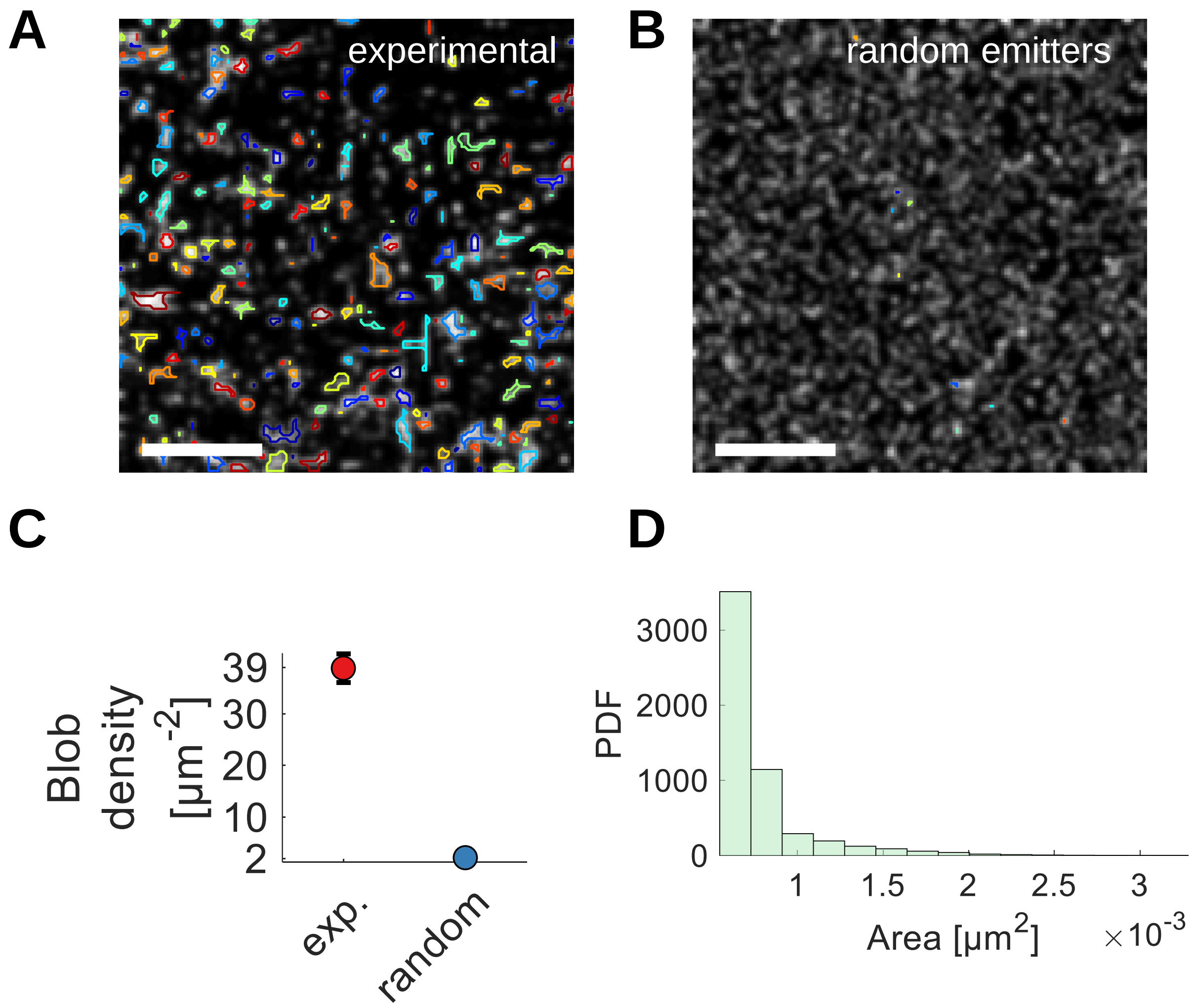


**Supplementary Figure 5: Segmentation on images of randomly distributed emitters. A)** An exemplary super-resolved image of chromatin *in vivo*, the identified blobs are overlaid and randomly colored. **B)** Emitters were randomly distributed and the segmentation algorithm was applied to those images. The number of emitter was matched to the number of beads of the modeled chromosomes within the imaging volume (compare Figure 2B). Scale bar is 0.5 µm. **C)** The blob density for blobs identified on experimental images and images containing randomly distributed emitters. Experimental images contained ~20-fold more blobs than could be identified using random images. **D)** Area distribution for blobs identified on images containing randomly distributed emitters (mean ± std.: $(\mathbf{0.4} \pm \mathbf{0.2}) \cdot \mathbf{10^{-3}}$ µm$^2$).

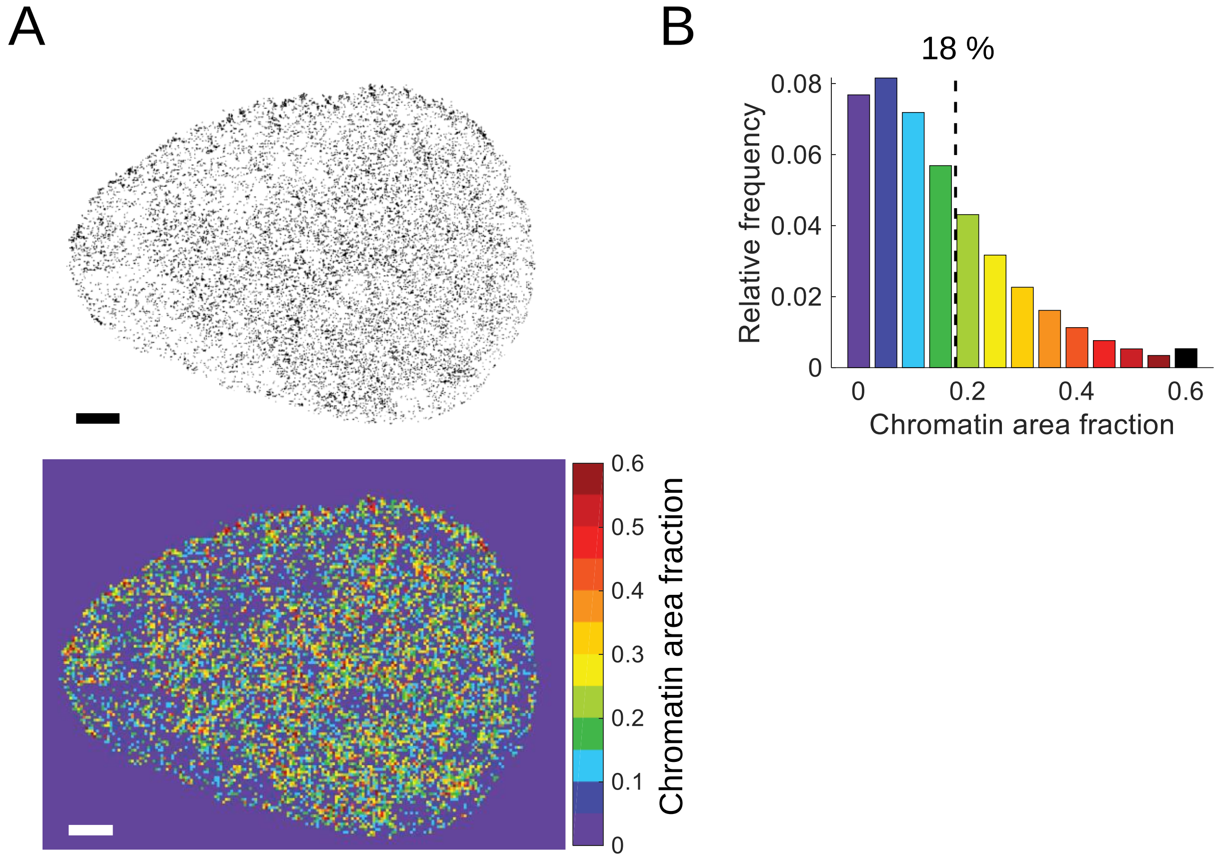


**Supplementary Figure 6: Chromatin area fraction. A)** Segmented images are divided into boxes with dimensions 120 nm x 120 nm and the chromatin area fraction is computed for each box. Exemplary, a map of chromatin area fractions is shown color-coded from low to high chromatin density (purple to red). Scale bar is 2 µm. **B)** Histogram of the observed chromatin area fractions. The black line denotes 18 ± 14 % (mean ± std). Data from 322 frames of two cells.

### SUPPLEMENTARY NOTE 2

#### Suitability of Optical Flow for super-resolution images of chromatin

Optical Flow is used to compute a flow field between two subsequent images in scenarios in which dynamic information cannot be retrieved from single-particle tracking approaches, for example, due to high labeling densities (*4*). Optical Flow algorithms are generally evaluated with respect to the angular error (AE), a measure for the error in the direction between a computed and a ground truth vector. Likewise, the endpoint error (EE) is a measure for the error in the magnitude. For two computed vectors $\boldsymbol{a}$ and $\boldsymbol{b}$, the AE and EE are computed as (*5*)

$$AE = \cos^{-1}\left(\frac{\boldsymbol{ab}}{|\mathbf{a}||\mathbf{b}|}\right) \tag{2}$$

and

$$EE = |\mathbf{a} - \mathbf{b}| \tag{3}$$

Current Optical Flow algorithms achieve sub-pixel EE and AE of around 20° for bulk chromatin imaging. Here, we prove that the Optical Flow algorithm used previously for conventional microscopes (*4*, *5*) results in comparable AE and EE values using super-resolved time series of chromatin. To this end, ground truth data is simulated as described previously (*4*). A density of 6 emitters per $\mu m^2$ and an acquisition time of 30 ms was used to resemble experimental data (Supplementary Figure 1). The images were either summed up directly in sets of 12 in order to achieve experimental time resolution of 360 ms or first processed by Deep-PALM and then summed. Optical Flow was computed for both sets and the AE and EE were computed. Exemplary simulated images and ground truth vectors, as well as computed flow fields, are superimposed in Supplementary Figure 7A-B. The resulting AE and EE from 20 independent runs are summarized in Supplementary Figure 7C. Optical Flow on super-resolved images did not significantly change the accuracy in direction, however the endpoint error is slightly smaller for flow fields computed on super-resolved images. These results thus validate Optical Flow for the use on super-resolution time series of chromatin.

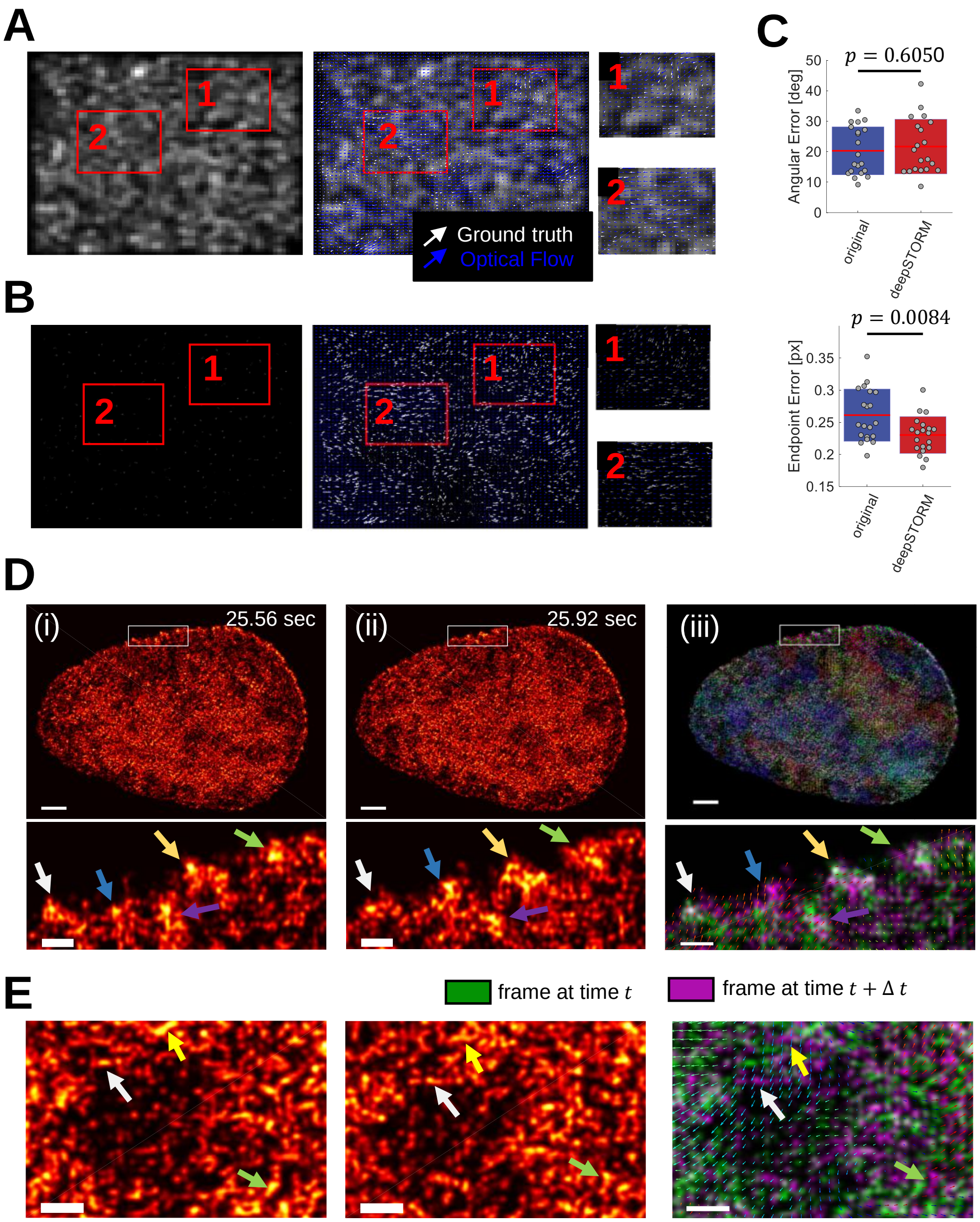


**Supplementary Figure 7: Performance of Optical Flow on conventional and super-resolved images and examples of super-resolution chromatin flow fields. A)** Exemplary simulated conventional fluorescence microscopy images (left), ground truth and estimated flow field (middle). Magnified regions as indicated by the red boxes (right). **B)** As A) for images analyzed with Deep-PALM. For visualization, only every eighth vector is shown (Deep-PALM images are up-sampled 8-fold compared to the input images). **C)** Angular and endpoint error over 20 independent sets of simulated images. Statistical significance was determined by a two-sided t-test. **D)** (i-ii) Two subsequent images of chromatin. Magnifications show prominent mobile blobs on the nuclear periphery in both images (colored arrows). (iii) The flow field and corresponding magnification are shown on top of a fused image of both super-resolved images in (i) and (ii) (green and purple respectively, co-localization are white). The flow field is colored according to the direction of vectors (see color wheel) **E)** As for D) in the nuclear interior. Scale bars: whole nucleus 2 µm, magnifications 500 nm. Flow vectors are not drawn to scale and down-sampled 8-fold for clarity.

**Supplementary Movie 1: Time series of super-resolved chromatin structure and dynamics.** The centroid positions of each identified blob are mapped onto the nucleus and colored according to their nearest-neighbor distance, area, mean displacement direction, and magnitude. Colors are given such that the respective maximum parameter value over the whole image series is red, the minimum parameter value is blue. Colors thus indicate parameter values relative to the parameter range. Displacement direction is color-coded according to the color-wheel shown.

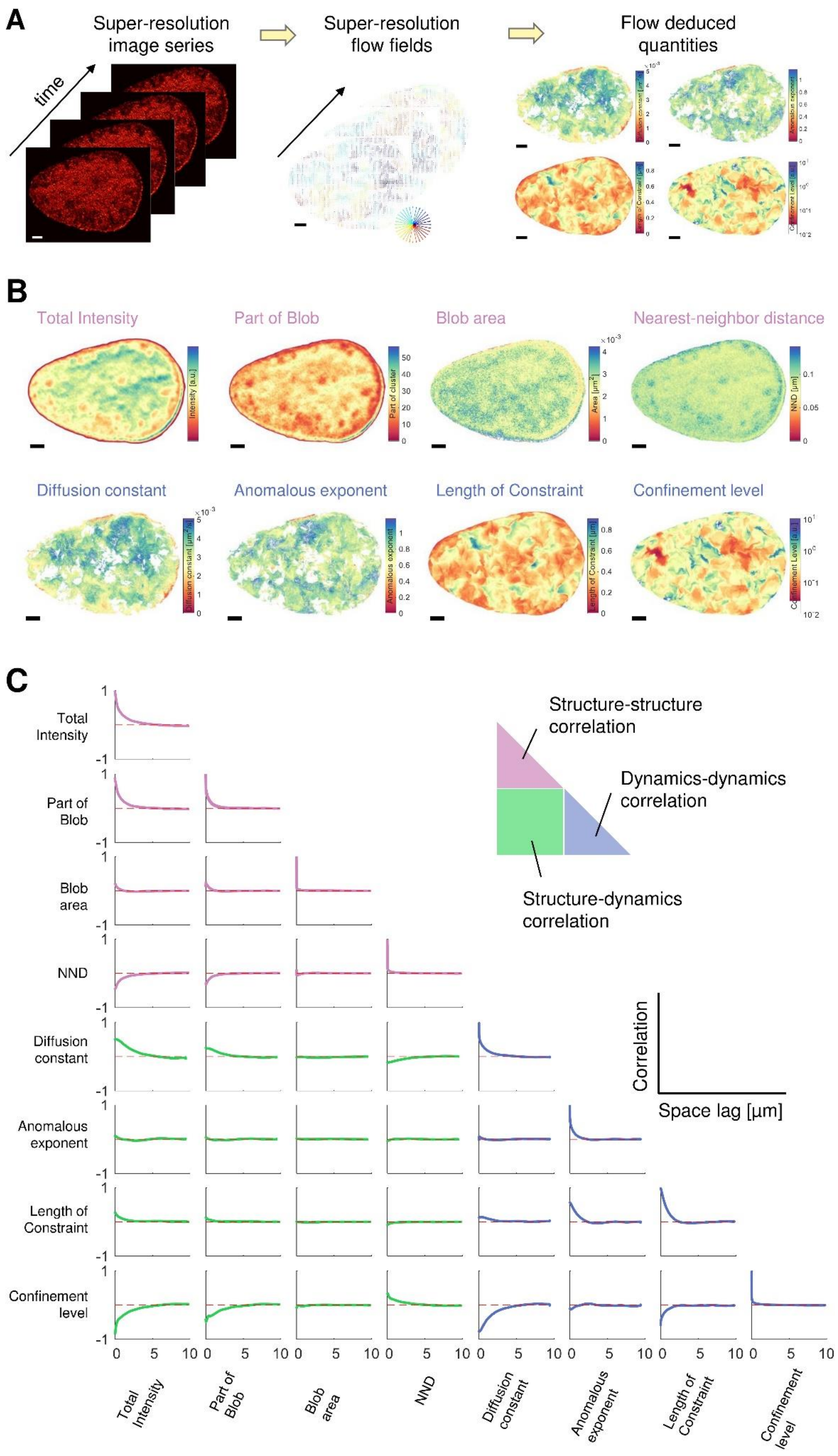
A
Super-resolution image series
time
Super-resolution flow fields
Flow deduced quantities
B
Total Intensity
Part of Blob
Blob area
Nearest-neighbor distance
Diffusion constant
Anomalous exponent
Length of Constraint
Confinement level
C
Total Intensity
Part of Blob
Blob area
NND
Diffusion constant
Anomalous exponent
Length of Constraint
Confinement level
Structure-structure correlation
Dynamics-dynamics correlation
Structure-dynamics correlation
Correlation
Space lag [μm]

**Supplementary Figure 8: Global spatial correlation of structural and dynamic parameters. A)** Illustration of the Hi-D workflow. A time series of super-resolution images (left panel) is input to the Optical Flow algorithm resulting in flow fields with a pixel size of 13.5 nm (middle panel). By trajectory reconstruction and motion classification, quantities describing the underlying bulk motion are computed. **B)** Eight parameters characterizing the global chromatin structure and dynamics during the whole time series are shown. The structural parameters (upper row) are the total intensity of super-resolved images, the counts how often each pixel was identified as part of a chromatin blob, the average blob area per pixel and the average nearest-neighbor distance for each pixel. Dynamic parameters are the diffusion constant and anomalous exponent, which was computed by regression of mean squared displacement curves (Materials and Methods), the length of constraint and the average confinement level. Scale bar is 3 µm. **C)** The spatial correlation between all combinations of structural and dynamic parameters over space lag is shown as average over two cells.

### SUPPLEMENTARY NOTE 3

#### t-SNE and its robustness with respect to distance metrics and perplexity

High-Dimensional parameter space is input to the t-SNE algorithm. The underlying principle is that data points, which are similar with respect to a number of factors (dimensions) lie close in high-dimensional space (employing a certain distance metric). The mapping into lower dimensionality (for instance in 2D) by t-SNE is initialized by assigning each point a random position in 2D (Figure 6). Illustratively, a set of springs between all data points exert a repelling or attractive force on each other depending on if the current distances between data points in 2D represent the distances between the data points in high-dimensional space. The 2D positions are iteratively refined in order to minimize the divergence between the high-dimensional and two-dimensional distributions.

More specifically, a high-dimensional pairwise distance measure can be defined between all points. The similarity of data points $\mathbf{x_i}$ and $\mathbf{x_j}$ is expressed as the conditional probability, $p_{j|i}$, that $\mathbf{x_i}$ would pick $\mathbf{x_j}$ as its neighbor under the assumption that neighbors are picked in proportion to their probability density under a Gaussian centered at $\mathbf{x_i}$ (*6*):

$$p_{j|i} = \frac{\exp\left(-\left\|\boldsymbol{x_i} - \boldsymbol{x_j}\right\|^2 / 2\sigma_i^2\right)}{\sum_{k \neq i} \exp(-\|\boldsymbol{x_i} - \boldsymbol{x_k}\|^2 / 2\sigma_i^2)}, \qquad (4)$$

and as the symmetrized conditional probabilities

$$p_{ij} = \frac{p_{j|i} + p_{i|j}}{2n}, \qquad (5)$$

where n is the number of data points. The variance of the Gaussian, $\sigma_i^2$, is determined by a binary search in order to obtain a user-specified value for the *Perplexity*, which is defined as

$$Perplexity(P_i) = 2^{H(P_i)}, \qquad (6)$$

where $H(P_i)$ is Shannon's entropy

$$H(P_i) = -\sum_j p_{j|i} \log_2 p_{j|i} \qquad (7)$$

and $P_i$ is the conditional probability distribution over all other data points given $\mathbf{x_i}$. The perplexity is, loosely speaking, controlling the number of close neighbors of each point and can have a complex non-linear influence of the resulting distribution of points.

The conditional probability $q_{j|i}$ of two points $\mathbf{y_i}$ and $\mathbf{y_j}$ in the two-dimensional space is modelled by a t-distribution:

$$q_{ij} = \frac{\left(1 + \left\|\boldsymbol{y_i} - \boldsymbol{y_j}\right\|^2\right)^{-1}}{\sum_{k \neq l}(1 + \|\boldsymbol{y_k} - \boldsymbol{y_l}\|^2)^{-1}} \qquad (8)$$

The algorithm first randomly assigns a position to each data point in two-dimensional space and then iteratively refines the position of data points such as to minimize the Kullback-Leibler (KL) divergence, a natural measure for the mismatch between the joint probability distributions in the high-dimensional space, P, and in the low-dimensional space, Q. Thus, the cost function at every iteration is the KL divergence between P and Q,

$$KL(P||Q) = \sum_i \sum_j p_{ij} \log\left(\frac{p_{ij}}{q_{ij}}\right), \tag{9}$$

which is minimized using a gradient descent method. In other words, the two-dimensional position $y_i$ is modified such as to minimize the KL divergence between P and Q. This minimization scheme depends critically on the conditional probabilities $p_{ij}$ and $q_{ij}$ and therefore on the distance $\|\cdot\|$ between points and the perplexity. We tested the influence of different distance metrics and perplexity values on our data set to exclude artifacts arising through an improper choice of parameters. The following distance metrics were tested:

1) Euclidian distance

$$\|\boldsymbol{x_i} - \boldsymbol{x_j}\| = \sqrt{(\boldsymbol{x_i} - \boldsymbol{x_j})^T (\boldsymbol{x_i} - \boldsymbol{x_j})} \tag{10}$$

2) Mahalanobis distance

$$\|\boldsymbol{x_i} - \boldsymbol{x_j}\| = \sqrt{(\boldsymbol{x_i} - \boldsymbol{x_j})^T S^{-1} (\boldsymbol{x_i} - \boldsymbol{x_j})} \tag{11}$$

where $S$ is the covariance matrix. The Mahalanobis distance reduces to the Euclidian distance if $S$ is the identity matrix.

3) Correlation distance

$$\|\boldsymbol{x_i} - \boldsymbol{x_j}\| = 1 - \frac{(\boldsymbol{x_i} - \boldsymbol{\bar{x}_i})^T (\boldsymbol{x_j} - \boldsymbol{\bar{x}_j})}{\sqrt{(\boldsymbol{x_i} - \boldsymbol{\bar{x}_i})^T (\boldsymbol{x_i} - \boldsymbol{\bar{x}_i})} \sqrt{(\boldsymbol{x_j} - \boldsymbol{\bar{x}_j})^T (\boldsymbol{x_j} - \boldsymbol{\bar{x}_j})}} \tag{12}$$

where $\boldsymbol{\bar{x}}$ denotes the average value of $\boldsymbol{x_i}$.

Perplexity values were varied from 30 to 200. Note that t-SNE is a probabilistic approach since points are initially distributed randomly in two dimensions. Therefore, multiple runs on the same data set might result in varying results.

Exemplary t-SNE maps are shown in Supplementary Figure 10A for the tested perplexity values and distance metrics. Maps are colored corresponding to Figure 6D. The embedding of points in two dimensions varies greatly among the scenarios. However, the probability of two points being nearest neighbors is largely conserved and thus ranking of input parameters yield similar results across the employed scenarios Supplementary Figure 10B). Rankings are especially robust when the distance metric is the Euclidian distance or the Mahalanobis distance. When the correlation distance is employed, the rankings slightly change. Especially, the area of blobs seems to be more prominent than the flow magnitude, in contrast to rankings when one of the other distance metrics is used. However, rankings change only to a small extent among the different distance metrics and perplexity values and the presented results are therefore free of artifacts of t-SNE or algorithm-dependent parameters.

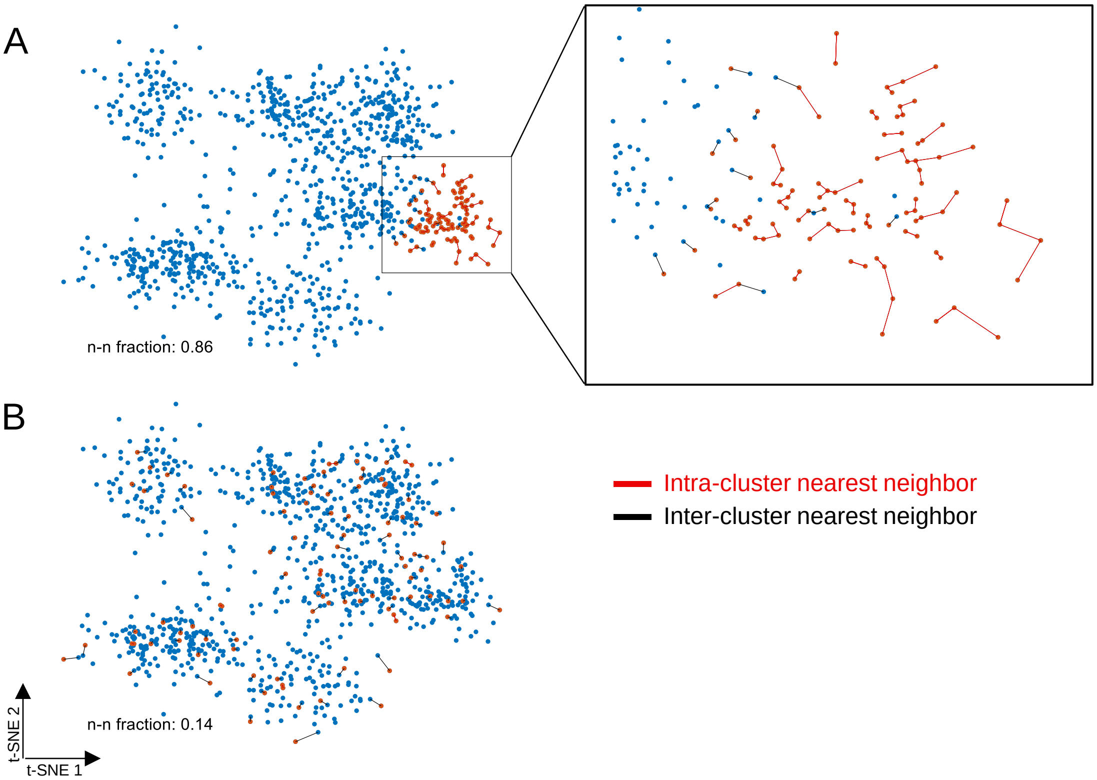


**Supplementary Figure 9: Clustering illustration of points within a subset based on nearest-neighbors in t-SNE maps.** Dimension reduction using t-SNE results in point clouds in which nearest neighbors in high-dimensional space are conserved and mapped as nearest neighbors in two dimensions. Points at which a number of interest exhibits unusually high/low values are identified (red points). Mapping of two points as nearest neighbors in reduced t-SNE space can be due to the fact that nearest neighbors belong to the same subset of high/low values of a variable of interest. To quantify this empirical characteristic, the number of nearest neighbors within the same subset in t-SNE space is counted relative to the total number of nearest neighbors of all points within the subset. **A)** A subset mostly contains its own nearest neighbors (86% of nearest neighbors of all points in the subset are contained within the subset). The magnification shows nearest neighbor connections within the subset (red lines) and between points in the subset and points not contained in the subset (black lines). **B)** Points within the subset are distributed over the whole t-SNE space and thus do not form a grouped region. The fraction of nearest neighbor links within the subset is small (14%) compared to the clustered case.

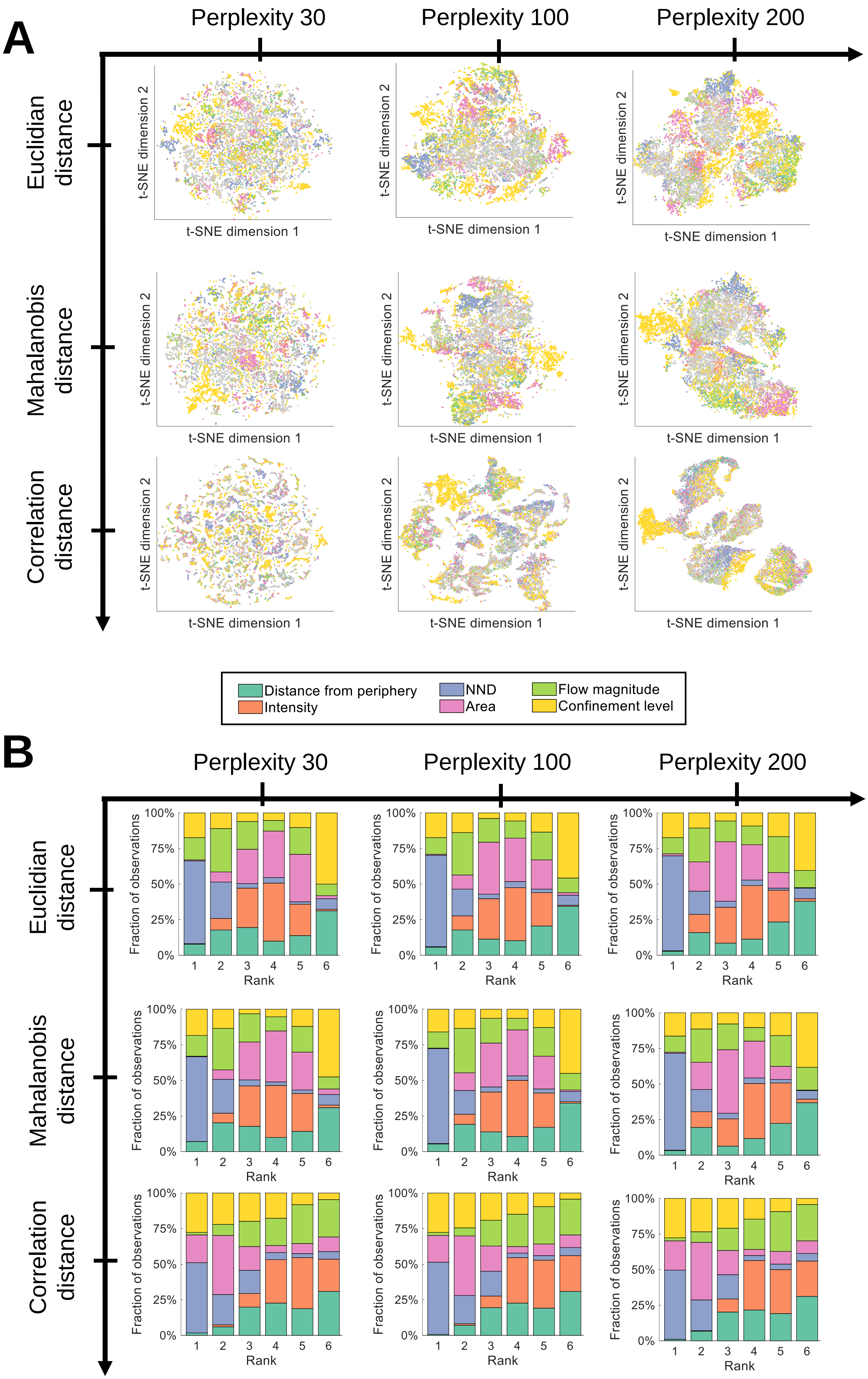
A
Perplexity 30
Perplexity 100
Perplexity 200
Euclidian distance
Mahalanobis distance
Correlation distance
t-SNE dimension 1
t-SNE dimension 2
Distance from periphery
Intensity
NND
Area
Flow magnitude
Confinement level
B
Fraction of observations
Rank
100%
75%
50%
25%
0%
1
2
3
4
5
6

**Supplementary Figure 10: t-SNE for different distance metrics and perplexity values.** We tested the influence of different distance measures between the high-dimensional data points as well as variations in the perplexity from 30 to 200 and found that our data are robust to changes in these parameters within the explored range. **A)** Exemplary t-SNE maps and **B)** feature ranking considering various distance metrics and values for the perplexity parameter applied to our data.
